# Donor-derived CD8^+^CD122^+^ Tregs generated in mixed donor chimeric NOD mice suppress autoreactive T cells

**DOI:** 10.64898/2026.03.20.712252

**Authors:** Shiva Pathak, Cameron S. Bader, Bettina P. Iliopoulou, Shobha Regmi, Pin-I Chen, Biki Gupta, Xiangni Wu, Blake Mosher, Alexandra Wells, Levon Witherspoon, Kayla Jenkins, William Harper, Emily SooHoo, Abigail Twoy, Rizwan Ahmed, Suparna Dutt, Nadine Nagy, Kent P. Jensen, Garrison Fathman, Avnesh S. Thakor, Mark M. Davis, Everett H. Meyer

## Abstract

The establishment of mixed hematopoietic chimerism is a promising way to induce immune tolerance for islet replacement therapy and to treat the underlying autoimmunity in type 1 diabetes (T1D). Mixed chimerism not only promotes effective thymic negative selection of autoreactive cells but also restores regulatory T cell (Treg) function and peripheral tolerance. In the current study, we determined that a novel class of donor-derived CD8^+^CD44^+^CD122^+^ Tregs (d-CD8^+^CD122^+^ Tregs) plays a crucial role in controlling autoimmunity in non-obese diabetic (NOD) mice with induced mixed chimerism. Using adoptive T cell transfer experiments, we showed that d-CD8^+^CD122^+^ Tregs abrogate autoimmunity by selectively depleting the exogenously injected diabetogenic T cells in Recombination-Activating Gene deficient NOD mice. These d-CD8^+^CD122^+^ Tregs from NOD chimeras show upregulation of Helios, Programmed cell death protein 1, perforin, granzyme-B, CD39, Folate receptor 4, and downregulation of proinflammatory markers like Scart1 and Scart2. Using *in vitro* assays, we show that d-CD8^+^CD122^+^ Tregs respond specifically to a Complementarity-Determining Region-3 peptide sequence derived from T cell receptors of islet antigen-specific autoreactive T cells. Similarly, we found that individuals with T1D have a deficiency in CD8^+^CD122^+^ Tregs, suggesting a potential loss of regulatory function accompanies disease onset. Revitalizing CD8^+^CD122^+^ Tregs may offer a new therapeutic strategy of restoring immune tolerance in autoimmune diabetes.

**One sentence summary:** Inducing mixed donor chimerism in NOD mice generates donor-derived CD8^+^CD122^+^ Tregs that suppress autoimmunity and restore immune tolerance by selectively eliminating autoreactive T cells.

## INTRODUCTION

Type 1 diabetes (T1D) is a chronic autoimmune disease characterized by insulin deficiency and hyperglycemia (1). The combined activities of antigen presenting cells (APCs) and autoreactive B and T cells result in the selective destruction of insulin-producing β cells in the islets of Langerhans (2, 3). In T1D, defects in thymic negative selection of developing T cells and reduced regulatory T cell (Treg) function in the periphery contribute to loss of immune tolerance to islet-derived self-antigens (4, 5). Tregs are essential for maintaining immune tolerance and act by preventing unchecked B and T cell immune responses. A highly studied ‘classical’ Treg population in T1D, CD4^+^CD25^+^Foxp3^+^ T cells, have been shown to be defective in function in both murine models and human subjects with T1D (6, 7). Loss of regulatory control from these cells permits infiltration and activation of islet antigen-specific effector T cells, thereby contributing to the autoimmune destruction of pancreatic islet β cells (6). In addition to the classical CD4^+^CD25^+^ Tregs, other immunoregulatory T cell subpopulations have been described and linked to T1D, including the recently identified CD8^+^CD44^+^CD122^+^ (CD8^+^CD122^+^) subset (8).

Jiang et al. demonstrated that CD8^+^ T cells function as a homeostatic regulatory compartment that limits the expansion of self-reactive CD4^+^ T cells in autoimmune encephalomyelitis (9). Although the CD8^+^ T cells have been implicated in the deletion of autoreactive CD4 T cell clones, the subset responsible for this activity remains unclear. CD8^+^CD122^+^ Tregs are a distinct CD8^+^ subset characterized by expression of CD122, the IL-2 receptor β-chain (10). These cells have been shown to suppress islet allograft rejection more effectively than conventional CD4^+^CD25^+^ Tregs, exhibiting greater suppressive activity and proliferative capacity (11). In NOD mice, however, CD8^+^CD122^+^ Tregs exhibit impaired suppressive functions compared to those from C57BL/6 mice, suggesting that functional defects within this subset may contribute to the loss of peripheral tolerance in autoimmune diabetes (8). Rifai et al. reported that CD8^+^CD122^+^ Tregs suppress activated T cells in an MHC class I-restricted manner, through conventional TCR-dependent recognition, whereas Qa-1, non-classical MHC class I, and MHC class II molecules are dispensable (12). More recently, a human CD8^+^ Treg subset expressing CD158 was identified as a critical regulator of autoimmunity in systemic lupus erythematosus. These CD8^+^CD158^+^ Tregs mediate suppression through direct contact-dependent killing of autoreactive CD4^+^ T cells *via* perforin- and granzyme B-mediated cytotoxicity. In mice, CD8^+^Ly49^+^ T cells are considered the functional and evolutionary counterpart of human CD8^+^CD158^+^ Tregs and have been proposed to constitute a component of the broader CD8^+^CD122^+^ regulatory compartment (13–15).

As a therapeutic strategy, we and others have demonstrated that autoimmune diabetes can be prevented in NOD mice through allogeneic hematopoietic stem cell transplantation (HSCT) that results in mixed donor chimerism. These studies have shown HSCT as a promising way to alleviate autoimmunity in murine T1D (5, 16, 17). Allogeneic HSCT helps restore immune homeostasis by correcting defects in central deletion and peripheral tolerance by generating functional donor-derived Tregs (18, 19). The latter mechanism may be important in this context as we and others have found that chimeric NOD recipients have a low-level persistence of circulating autoreactive T cells. Nevertheless, mixed chimeric NOD mice are protected from diabetes onset (17). However, the precise mechanism by which mixed donor chimerism prevents T1D onset remains poorly understood. Thus, mixed chimerism in NOD mice represents an ideal therapeutic system to evaluate how peripheral tolerance pathways can be restored in autoimmune diabetes.

Here, we revealed the critical involvement of donor-derived CD8^+^CD122^+^ (d-CD8^+^CD122^+^) Tregs in the maintenance of peripheral tolerance after HSCT. We demonstrated that d-CD8^+^CD122^+^ Tregs were sufficient to induce peripheral tolerance in the Recombination-Activating Gene 1 deficient NOD (NRG) recipients. We performed comprehensive flow cytometric and transcriptomics analyses to identify the unique gene and protein signatures of the tolerogenic d-CD8^+^CD122^+^ Tregs. We showed that d-CD8^+^CD122^+^ Tregs are capable of selectively depleting islet β-antigen specific CD8^+^ T cells through the recognition of Complementarity Determining Region 3 (CDR3)-derived peptides of islet-specific glucose-6-phosphatase catalytic subunit-related protein (IGRP)-reactive CD8^+^ T cells. Using bulk RNA-Seq and confirmatory flow cytometry (protein) analyses, we found that CD8^+^CD122^+^ Tregs are significantly diminished in diabetes-prone NOD mice and in humans with T1D. The NOD CD8^+^CD122^+^ Tregs exhibit phenotypic and functional defects with regards to expression of immune regulatory surface markers, impaired metabolic pathways and suppressive capacity. CD8^+^CD122^+^ Tregs from patients with T1D also exhibit deficiencies in number, function, and gene signatures. These findings support potential therapeutic strategies that induce or directly use CD8^+^CD122^+^ Tregs to correct the functional deficiencies observed in T1D patients and reveal new details that support mixed hematopoietic chimerism for the prevention of autoimmunity.

## RESULTS

### Non-obese diabetic mice have defective CD8^+^CD122^+^ Tregs

To explore the phenotypic and functional properties of CD8^+^CD122^+^ Tregs in NOD mice, we conducted a comprehensive phenotyping profiling and functional assays, comparing them with age-matched female non-obese diabetes-resistant (NOR) CD8^+^CD122^+^ Tregs. NOD and NOR mice were euthanized at the age of 14 weeks and CD8^+^CD122^+^ Tregs were sorted for suppression assay and bulk RNA sequencing. NOR mice were used for comparison since they are syngeneic in major histocompatibility complex (MHC) to the NOD mice, with a key difference that NOD mice spontaneously develop diabetes while the NOR mice do not (**Fig 1a-b, S1**). Results showed that NOD mice have a significantly lower percentage of CD8^+^CD122^+^ Tregs compared to NOR mice in peripheral blood (6.2% versus 11.9%; *p*=0.0005) (**Fig 1c**). There were no statistically significant differences in multiple surface markers tested, except for increased CD38 expression in NOD CD8^+^CD122^+^ Tregs (**Fig S2**). We then evaluated suppressive function of NOD versus NOR-derived CD8^+^CD122^+^ Tregs using a standard suppression assay. We cocultured CD8^+^CD122^+^ Tregs with cell trace violet (CTV)-labeled third-party (FVB mice) T conventional cells and CD3/CD28 stimulation beads in RPMI-1640 media for 5 days (**Fig 1d**). The use of third-party responder T cells reduces donor-specific variability and allows evaluation of Treg function across allo-contexts (20). Representative flow cytometry plots showing CTV dilution in T conventional cells are depicted in **Fig 1e**. Percentages of T cell suppression at different ratios of T conventional cells and CD8^+^CD122^+^ Tregs revealed impaired effector function of the CD8^+^CD122^+^ Tregs in NOD mice compared to NOR (*p*<0.01), suggesting that suppressive function defects among CD8^+^CD122^+^ Tregs may contribute to loss of peripheral tolerance and disease pathogenesis in NOD mice.

**Fig 1.**
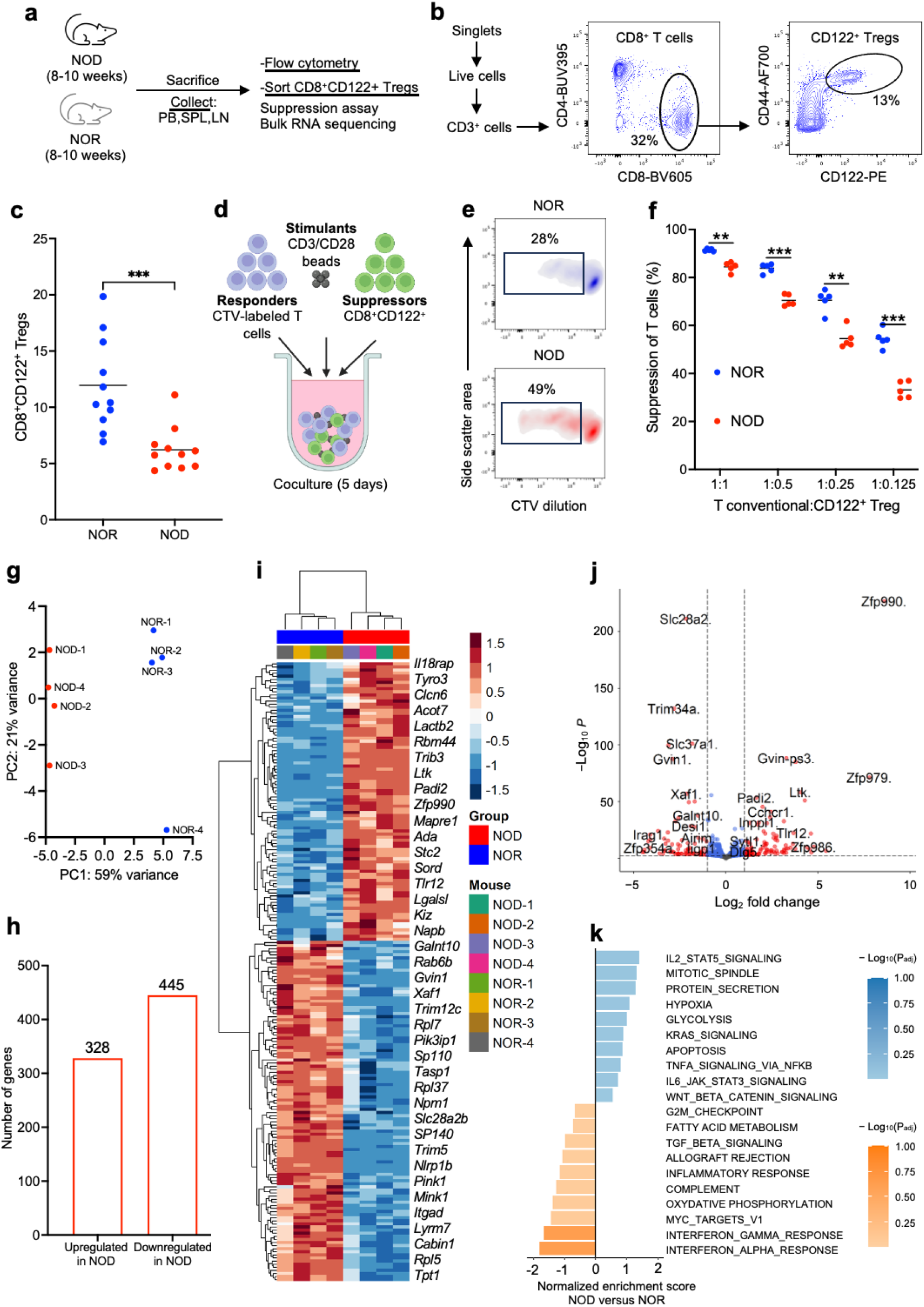
NOD mice have defective CD8^+^CD122^+^ Tregs. (**a**) Overall experiment design to evaluate phenotype and function of NOD CD8^+^CD122^+^ Tregs. (**b**) Gating hierarchy of CD8^+^CD122^+^ Tregs from singlet live CD3^+^ T cells. (**c**) Frequency of CD8^+^CD122^+^ Tregs in peripheral blood of age- and sex-matched NOR and NOD mice. (**d**) Design of *in vitro* suppression assay. CTV-labeled third-party (FVB) responder T cells were co-cultured with CD8^+^CD122^+^ Tregs in the presence of CD3/CD28 activation beads for 5 days. (**e**) Representative density plots showing dilution of CTV in responder T conventional cells. (**f**) Percent suppression of T conventional cells at different ratios of T conventional cells and CD8^+^CD122^+^ Tregs. Data pooled from two independent experiments. (**g-k**) Bulk RNA sequencing of NOD CD8^+^CD122^+^ Tregs comparing with age- and sex-matched NOR CD8^+^CD122^+^ Tregs. (**g**) Principal component (PC) analysis of gene expression from NOD CD8^+^CD122^+^ Tregs and NOR CD8^+^CD122^+^ Tregs. (**h**) Total number of upregulated and downregulated genes in NOD CD8^+^CD122^+^ Tregs compared to NOR CD8^+^CD122^+^ Tregs. (**i**) Heatmap and hierarchical clustering of top 200 most differentially expressed genes by adjusted *p*-value in NOD CD8^+^CD122^+^ Tregs *versus* NOR CD8^+^CD122^+^ Tregs. (**j**) Volcano plot showing gene expression log_2_ fold change and –log_10_(P_adj_) in NOD CD8^+^CD122^+^ Tregs *versus* NOR CD8^+^CD122^+^ Tregs. Significant differentially expressed genes are marked in red and are defined by a log_2_ fold change > |1| and a P_adj_ < 0.001. (**k**) Gene-set enrichment analysis of NOD CD8^+^CD122^+^ Tregs *versus* NOR CD8^+^CD122^+^ Tregs. The 20 most immune-related and differentially expressed pathways by normalized enrichment score are shown. \*\**p*<0.01, \*\*\**p*<0.001. CTV; cell trace violet, NOD; non-obese diabetic, NOR; non-obese diabetes-resistant.

To further explore the differences between NOD and NOR CD8^+^CD122^+^ Tregs, we sorted CD8^+^CD122^+^ Tregs from NOD and NOR mice and performed bulk RNA sequencing. A detailed analysis revealed differences in gene expression among sorted NOD and NOR CD8^+^CD122^+^ Tregs. Principal component analysis revealed similarities among the samples within a group and showed a significant variance in the gene profile between the CD8^+^CD122^+^ Tregs of NOD and NOR mice (**Fig 1g**). A differential gene analysis identified a total of 328 upregulated and 445 downregulated genes (**Fig 1h**) in NOD *versus* NOR mice. Upregulation of *Il18rap* expression in NOD CD8^+^CD122^+^ Tregs suggests to a proinflammatory shift and compromised regulatory stability, consistent with dysregulated suppressive function and a compensatory IL-2/STAT5 signaling response to maintain suppressive capacity under inflammatory conditions. Unexpectedly, *Tlr12* transcripts were detected in NOD CD8^+^CD122^+^ Tregs from NOD mice. Given that these cells exhibited a pronounced pro-inflammatory transcriptional profile compared to NOR CD8^+^CD122^+^ Tregs, we interpret *Tlr12* expression as part of an inflammation-associated, innate-like transcriptional program. Another possibility is that increased mRNA abundance does not translate into detectable surface protein expression, potentially because of post-translational regulation, inefficient translation, or altered protein processing and trafficking. Alternatively, these transcripts may have regulatory functions that remain to be defined. NOD CD8^+^CD122^+^ Tregs exhibited downregulation of a broad set of genes critical for immune cell activation and effector function, a pattern associated with attenuated IFN-α and IFN-γ signaling. This transcriptional profile may reflect intrinsic defects in the CD8^+^CD122^+^ Tregs or shift in cytokine milieu that disrupt IFN-driven cross-talk with APCs, natural killer cells, and effector cells including autoreactive T cells. Moreover, the concurrent downregulation of *Slc37a1* and *Slc28a2b* together with reduced expression of key ribosomal genes (*Rpl7*, *Rpl37*, *Rpl9*, *Rpl18*, *Rpl17*, *Rpl32, Rpl5)* suggests a coordinated impairment of metabolic and protein synthesis pathways in NOD CD8^+^CD122^+^ Tregs, potentially compromising their suppressive function (**Fig 1i-k**).

### HSCT corrects functional deficiency of CD8^+^CD122^+^ Tregs

Based on our findings that revealed functional deficiencies in NOD CD8^+^CD122^+^ Tregs, we hypothesized that HSCT to induce mixed hematopoietic chimerism may overcome the observed deficiencies. A schematic representation of immune conditioning regimen is depicted in **Fig 2a**. CD45.1 NOD mice were preconditioned with a non-myeloablative conditioning comprising 8 doses of total lymphoid irradiation (TLI), 2 low doses of total body irradiation (TBI), and 5 doses of anti-thymocyte serum (ATS) prior to allogenic BMT from CD45.2 C57BL/6 donors (**Fig S3**), as reported previously by our group (17). Donor engraftment in blood was measured by flow cytometry *via* the presence of CD45.2^+^ immune cells in transplanted mice and was used to quantify percent mixed donor chimerism (**Fig 2b**). Whole-blood and T cell chimerism were found to be approximately 80% and 60% at day 60 post-transplant, respectively (**Fig 2c**). The induction of mixed chimerism led to an increase in the frequency of CD8^+^CD122^+^ Tregs in the chimeric NOD (9.2%) compared to the WT NOD mice (6.1%), suggesting that a transient lymphodepletion followed by HSCT increased the peripheral CD8^+^CD122^+^ Treg numbers (**Fig 2d**).

**Fig 2.**
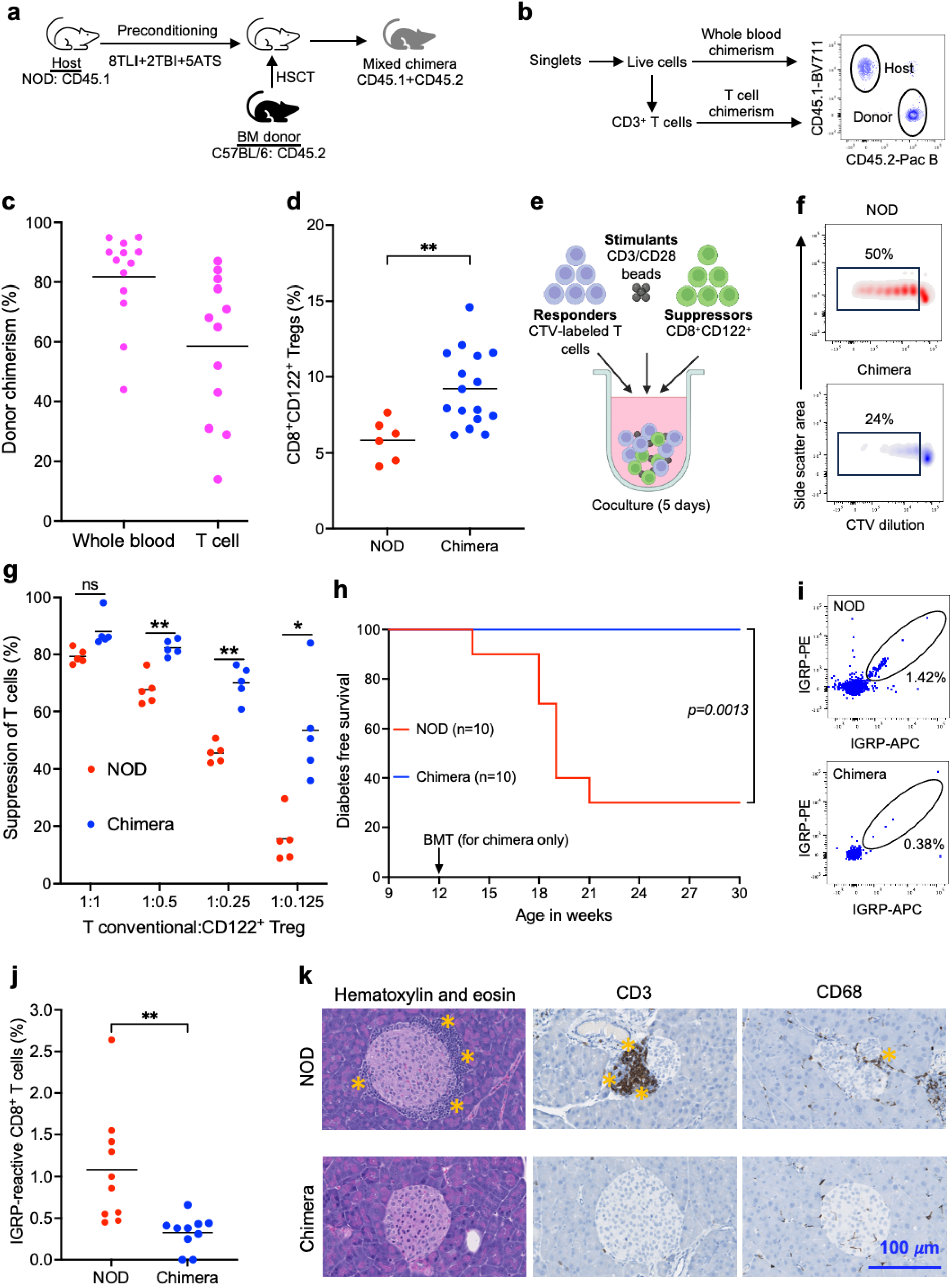
HSCT corrects functional deficiency of CD8^+^CD122^+^ Tregs. (**a**) Experiment design. Prediabetic NOD mice (10 weeks old) were preconditioned with 8 doses of TLI (2.4 Gy each), 2 doses of TBI (1.5 Gy each), and 5 doses of ATS (50 μL each). Allogeneic BM derived from C57BL/6 mice (50 × 10^6^) was transplanted into the preconditioned mice by intravenous injection. (**b**) Mice were bled at day 60 post-HSCT for mixed donor chimerism evaluation by flow cytometry. Host cells were labeled with CD45.1 and donor cells were stained with CD45.2 antibodies. (**c**) Evaluation of whole blood and T cell chimerism. (**d**) Frequency of CD8^+^CD122^+^ Tregs in age- and sex-matched WT NOD and chimeric mice at day 60 post-HSCT. (**e**) Design of *in vitro* suppression assay to evaluate the functionality of chimeric CD8^+^CD122^+^ Tregs. (**f**) Representative density plots showing dilution of CTV in T conventional cells after co-culture with CD8^+^CD122^+^ Tregs for 5 days. (**g**) Percent suppression of T conventional cells at different ratios of T conventional cells and CD8^+^CD122^+^ Tregs. Data pooled from two independent experiments. (**h**) Kaplan-Meier curve showing diabetes-free survival of WT NOD and chimera. Non-fasting blood glucose was monitored until week 30 post-HSCT and mice showing two consecutive glucose readings over 300 mg/dL were considered diabetic. (**i**) Representative flow cytometry plots showing IGRP-reactive CD8^+^ T cell staining in WT NOD and chimera. (**j**) Frequency of IGRP-reactive CD8^+^ T cells were measured in the peripheral blood of chimeric mice at day 60 post-HSCT. (**k**) Representative data showing hematoxylin and eosin, CD3, and CD68 staining of pancreas section obtained from WT NOD and chimeric NOD mice at day 60 post-HSCT. Asterisks indicate areas of infiltration. Magnification: 40×. Scale bar: 100 *μ*m. \**p*<0.05, \*\**p*<0.01, \*\*\**p*<0.001, \*\*\*\**p*<0.0001. ATS; anti-thymocyte serum, CTV; cell trace violet, HSCT; hematopoietic stem cell transplantation, IGRP; islet-specific glucose-6-phosphatase catalytic subunit-related protein, NOD; non-obese diabetic, TBI; total body irradiation, TLI; total lymphoid irradiation, WT; wildtype.

We then tested whether the defective suppressive function of NOD CD8^+^CD122^+^ Tregs could also be corrected following HSCT and the establishment of mixed chimerism. We again utilized a suppression assay (**Fig 2e**) in which CD8^+^CD122^+^ Tregs from mixed chimeric or WT NOD mice were sorted and co-cultured with third-party (FVB) T cells for 5 days in the presence of CD3/CD28 stimulation beads and proliferation of T cells was quantified. Representative flow cytometry plots in **Fig 2f** show dilution of CTV as a measure of proliferation of the third-party T conventional cells. We observed a statistically significant reduction in proliferation of the responder T cells in coculture with CD8^+^CD122^+^ Tregs from mixed chimera compared to those from WT NOD mice (*p*<0.05) (**Fig 2g**). This restoration of suppressive function of CD8^+^CD122^+^ Tregs following HSCT and establishment of mixed chimerism suggests that donor-derived CD8^+^CD122^+^ Tregs may be useful to abrogate autoimmunity in diabetes-prone NOD mice.

To assess durability of disease prevention, we monitored the chimeric mice until 30 weeks of age and there was no progression to diabetes observed **(Fig 2h)**. As expected, the WT NOD control mice developed hyperglycemia at age 14 weeks whereby over 70% of the control mice were diabetic by week 21. We next evaluated autoreactive T cells in the WT NOD and chimeric mice by flow cytometric staining using IGRP peptide loaded H-2K^d^, APC and PE-labeled tetramers (**Fig 2i, S4a**). At day 60 post-HSCT, the percentage of IGRP-reactive CD8^+^ T cells in the peripheral blood of HSC transplanted, mixed chimeric mice were significantly reduced compared to the WT NOD mice (0.3% *versus* 1.0%; *p*<0.01) (**Fig 2j**). Although fewer autoreactive T cells were present in chimeric mice, their phenotype was indistinguishable from WT NOD autoreactive T cells, implying functional equivalence (**Fig S4b**). Furthermore, hematoxylin and eosin staining of pancreas sections demonstrated a massive peri-islet infiltration in the pancreata of age and sex-matched WT NOD mice, but only minimal peri-islet infiltration in the chimeric mice at day 60 post-HSCT. Further immunohistochemistry staining with CD3 and CD68 revealed a significant presence of T cells and macrophages, respectively, in WT NOD mice. In contrast, the pancreata of chimeric mice showed minimal infiltration of both T cells and macrophages (**Fig 2k**). Notably, flow cytometry data demonstrated that the pancreatic T-cell repertoire in chimeric mice is largely donor-derived, with 100% of CD8^+^CD122^+^ Tregs deriving from the donor, indicating complete replacement of the defective pancreatic CD45.1^+^CD8^+^CD122^+^ Tregs (**Fig S5**).

### CD4^+^CD25^+^ Tregs are not required to maintain peripheral tolerance in chimeric mice

Despite the detection of low levels of IGRP-reactive CD8^+^ T cells in the chimeric mice, there was no incidence of hyperglycemia or histological evidence of overt insulitis. These data favor that there is a restoration of peripheral tolerance by the Tregs generated in the NOD chimera. To explore the role of the different Treg subtypes in the chimeric mice, we performed adoptive T cell transfer into NOD-Rag1^null^IL2rg^null^ (NRG) mice using T cells derived from the chimeric mice. The advantage of this approach is that by selecting different T regulatory cell subsets for adoptive transfer, it is possible to identify which subsets are necessary or sufficient for protection against insulitis. We sacrificed the chimeric mice 60 days after HSCT and harvested the spleen and lymph nodes to isolate T cells (**Fig 3a**). The sorting strategy and gating hierarchy are shown in **Fig 3b**. Donor and host-derived T cells were distinguished based on CD45.2 and CD45.1 staining, respectively. **Fig 3c** outlines the experimental design for the adoptive T cell transfer studies. Each NRG recipient received 2 × 10^6^ T cells. Mice that received CD3^+^ T cells isolated from wild-type (WT) NOD mice served as the control group. To evaluate the contribution of recipient-derived T cells, NRG mice were transferred with either CD45.1^+^CD3^+^ T cells or CD25-depleted CD45.1^+^CD3^+^ T cells. To assess the combined effects of donor- and host-derived T cells, recipient mice received either total CD3^+^ T cells isolated from mixed chimeras or CD25-depleted CD3^+^ T cells from mixed chimeras. Data showed that the adoptively transferred T cells survived and persisted in the NRG mice. The percentage of T cells at day 60 post-transplant in all groups receiving T cells from chimeric mice was lower while compared to the group that received WT NOD T cells (*p*<0.05), indicating slower expansion of chimera-derived T cells compared to that of the WT NOD T cells. However, there was no statistically significant difference among all other groups that received T cells from the chimeric mice (**Fig 3d**).

**Fig 3.**
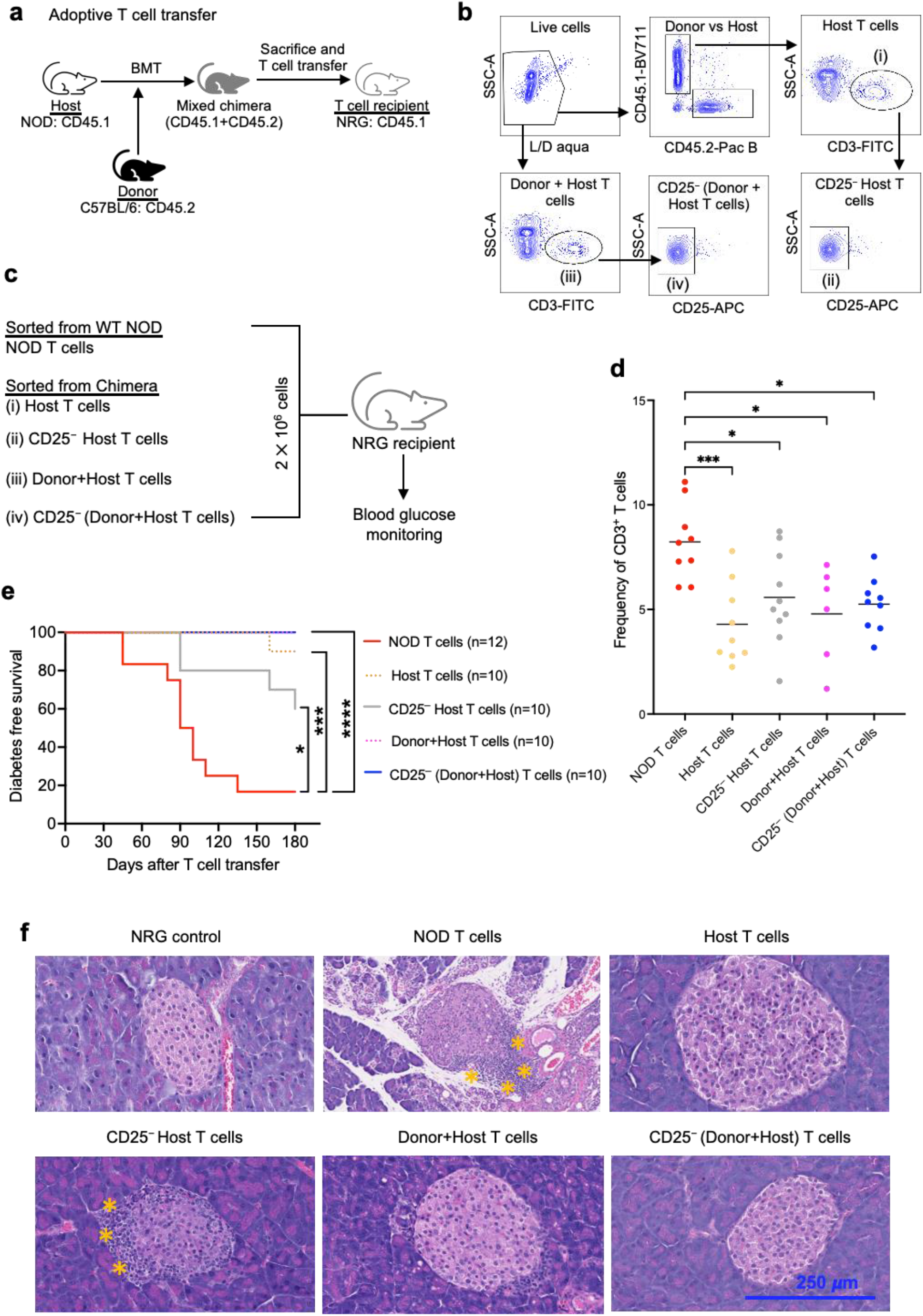
CD4^+^CD25^+^ Tregs are not required to maintain peripheral tolerance in chimeric mice. (**a**) Design of adoptive T cell transfer experiment. Chimeric mice were generated using C57BL/6 as donors and NOD as recipients. The chimeric mice were sacrificed at day 60 post-HSCT to isolate T cells by sorting. Adoptive T cell transfer was performed in NRG mice. (**b**) Gating hierarchy for sorting T cells from NOD chimera. Total T cells with or without CD25^+^ T cells were sorted from CD3^+^ fraction of the live cells while host-derived (NOD-derived) T cells were sorted from CD3^+^ fraction of live CD45.1^+^ cells. (**c**) List of all groups used in adoptive T cell transfer experiment. CD3^+^ T cells from WT NOD mice were sorted for transfer into the control group. (**d**) Persistence of T cells in peripheral blood of NRG mice after adoptive T cell transfer. (**e**) Kaplan-Meier curve showing diabetes-free survival of NRG mice after adoptive T cell transfer. Data were pooled from two independent experiments. (**f**) Representative data showing hematoxylin and eosin staining of pancreas section obtained from different groups. Asterisks indicate areas of infiltration. Magnification: 40×. Scale bar: 250 *μ*m. \**p*<0.05, \*\**p*<0.01, \*\*\**p*<0.001, \*\*\*\**p*<0.0001. HSCT; hematopoietic stem cell transplantation, NOD; non-obese diabetic, NRG; NOD-Rag1^null^IL2rg^null^, WT; wildtype.

Blood glucose levels of NRG mice were then monitored for 180 days following adoptive T cell transfer. The overall diabetes-free survival of the NRG mice is shown in **Fig 3e**. As expected, the NRG mice that received T cells from WT NOD mice became hyperglycemic (80%) by day 180 post adoptive transfer. When we transferred only host-derived T cells from the NOD chimera, diabetes was prevented with a low incidence of hyperglycemia (10%) at day 180 post T cell transfer. Interestingly, when CD25^+^ T cell depleted host-type T cells (CD25^−^ host T cells) were transferred into the NRG mice, 40% of recipient NRG mice became hyperglycemic indicating that adoptively transferred T cells are unable to effectively maintain peripheral tolerance in the absence of host-derived CD25^+^ T cells. Our data further demonstrated that host-derived CD4^+^CD25^+^ T cells from NOD chimeras exhibited increased PD-1 expression and enhanced suppressive function compared to CD4^+^CD25^+^ T cells from WT NOD mice. (**Fig S6**). These findings are consistent with previous studies showing that TLI/ATS conditioning upregulates PD-1 expression on host CD4^+^CD25^+^ T cells, thereby enhancing their suppressive function in the Balb/c-to-C57BL/6 HSCT model (21). Moreover, the transfer of total T cells (Donor+Host T cells) from the NOD chimera prevented the onset of hyperglycemia and diabetes in the recipient NRG mice (0% incidence), indicating that HSCT established immune homeostasis and maintained peripheral tolerance in NOD mice. Surprisingly, when the CD25^+^ T cells were depleted from total T cells that have both donor and host-derived T cells [CD25^−^ (donor+host) T cells] and adoptively transferred, none of the NRG recipients developed diabetes. This suggested an involvement of donor-derived non-CD25^+^ Tregs that can suppress autoimmunity and maintain a robust peripheral tolerance in the NRG recipients (**Fig 3e**).

We performed hematoxylin and eosin staining of the pancreas at day 60 post T cell transfer to determine if there was any leukocyte infiltration into the islets. As expected, immune cells were barely detected in the NRG controls which did not receive T cells, indicating absence of autoimmunity in these mice. In contrast, the pancreas of NRG mice that received WT NOD T cells had fewer islets overall and showed a massive immune cell infiltration in the peri-islet region indicating an overt autoimmunity. In mice that received host-derived T cells from chimeric donors, minimal immune cell infiltration was observed. Conversely, significantly higher immune cell infiltrates were detected following adoptive transfer of CD25^−^ host T cells. Furthermore, in the group that received donor+host T cells from chimeric mice, no immune cell infiltration was observed. Notably, CD25^−^ donor+host T cell group showed a minimal infiltration of immune cells in the pancreas (**Fig 3f**). These data further support the notion that donor-derived Tregs that lack CD25 expression are involved in the disease protection in NRG recipients.

### Donor CD8^+^CD122^+^ Treg depletion prior to adoptive T cell transfer leads to autoimmune diabetes in NRG mice

Given our findings that CD8^+^CD122^+^ Tregs are altered and appear dysfunctional in NOD mice, we hypothesized that d-CD8^+^CD122^+^ Tregs generated in the chimeric NOD recipients suppress autoreactive T cell expansion and diabetes in NRG mice. We euthanized NOD chimeras at day 60 post-HSCT, sorted CD8^+^ T cells, and performed adoptive T cell transfer experiments into NRG mice (**Fig 4a-c**). For the adoptive T cell transfer, donor-derived CD8^+^ T cells isolated from NOD chimera were identified as CD45.2^+^CD8^+^ T cells (**Fig 4b**). We also performed a simultaneous co-transfer of diabetogenic T cells from 14 weeks old female NOD mice into the CD8^+^ T cell receiving NRG mice (**Fig 4c**). Given the delayed diabetes onset in NRG mice by CD25^+^ T cells in our earlier adoptive T cell transfer experiments (**Fig 3**), the NRG recipients in this experiment received 3 × 10^6^ CD25^+^ T cell-depleted WT NOD T cells to accelerate disease onset. Over 80% of the NRG mice receiving WT NOD T cells became hyperglycemic by day 60 post-transplant. The transfer of 2 × 10^6^ CD8^+^ T cells from WT C57BL/6 mice did not prevent the disease onset. Importantly, diabetes onset was prevented in NRG mice when 2 × 10^6^ CD45.2^+^CD8^+^ T cells from NOD chimera were given at the time of WT NOD T cell injection. When CD122^+^ subset was depleted from the CD45.2^+^CD8^+^ T cells of NOD chimera [CD45.2^+^CD8^+^ T cells (CD122^dep^)] before the transplant, the protection was lost and the NRG recipients rapidly developed diabetes (**Fig 4d**). Flow cytometric analysis confirmed deletion of IGRP-reactive CD8^+^ T cells in the pancreas of mice that received CD45.2^+^CD8^+^ T cells, but the effect was lost when the CD122^+^ subset was depleted (**Fig 4e**). These data show that d-CD8^+^CD122^+^ Tregs eliminate autoreactive T cells and prevent diabetes onset.

**Fig 4.**
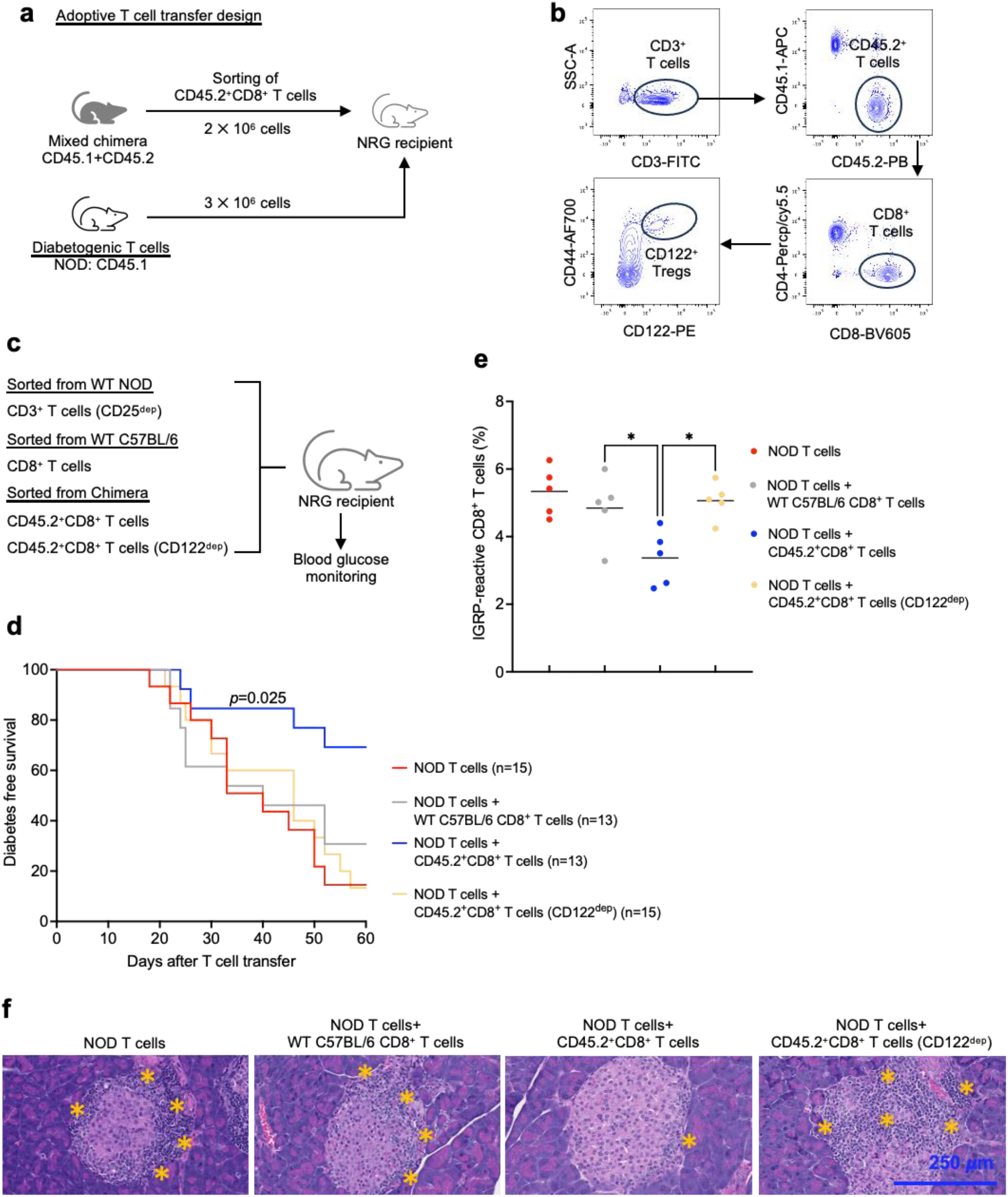
Donor CD8^+^CD122^+^ Treg depletion prior to adoptive transfer leads to autoimmune diabetes in NRG mice. (**a**) Design of adoptive T cell transfer experiment. Chimeric NOD mice were generated by transplanting allogeneic BM (50 × 10^6^) from C57BL/6 mice into the preconditioned NOD mice. NOD chimeras were sacrificed at day 60 post-HSCT for T cell sorting. Diabetogenic T cells from WT NOD mice (3 × 10^6^) and sorted CD45.2^+^CD8^+^ T cells (2 × 10^6^) ± CD8^+^CD122^+^ Tregs were co-transferred into NRG mice, and the mice were monitored for diabetes onset. (**b**) Gating hierarchy for T cell sorting for adoptive T cell transfer. CD3^+^ T cells were isolated and gated on CD45.2 positivity to sort d-CD8^+^CD122^+^ Tregs from NOD chimera. (**c**) Experimental groups. Diabetogenic CD3^+^ T cells were sorted from 12 weeks old prediabetic WT NOD mice. C57BL/6 mice that were age- and sex-matched to the BM donors were used to sort WT CD8^+^ T cells. From chimeric NOD mice, CD45.2^+^CD8^+^ T cells (donor CD8^+^ T cells) or CD122-depleted donor CD8^+^ T cells [CD45.2^+^CD8^+^ T cells (CD122^dep^)] were sorted. (**d**) Kaplan-Meier curve showing diabetes-free survival of NRG mice after adoptive T cell transfer. T cell recipient NRG mice were monitored until day 60 post T cell transfer. Data were pooled from three independent experiments. (**e**) Frequency of IGRP-reactive CD8^+^ T cells in pancreas of NRG recipients at day 60 post T cell transfer. NRG recipients were sacrificed, pancreas were digested using collagenase, and lymphocytes were isolated by density gradient centrifugation using Lympholyte^®^. (**f**) Representative data showing hematoxylin and eosin staining of pancreas section obtained from different groups. Magnification: 40×. Scale bar: 250 *μ*m. \**p*<0.05. BM; bone marrow, CD122^+^ Tregs; CD8^+^CD122^+^ Tregs, HSCT; hematopoietic stem cell transplantation, IGRP; islet-specific glucose-6-phosphatase catalytic subunit-related protein, NOD; non-obese diabetic, NRG; NOD-Rag1^null^IL2rg^null^, WT; wildtype.

To further confirm that the d-CD8^+^CD122^+^ Tregs protect from peri-islet lymphocyte infiltration and autoimmune diabetes, we performed hematoxylin and eosin staining of pancreas sections of recipient NRG mice given diabetogenic T cells and CD8^+^ T cells (**Fig 4f**). NRG mice given NOD T cells only or NOD T cells + WT C57BL/6 CD8^+^ T cell transfer showed a massive peri-islet immune cell infiltration. In contrast, minimal infiltration was observed in the pancreas sections of the NRG mice reconstituted with NOD T cells + CD45.2^+^CD8^+^ T cells from NOD chimera. This was expected given the fact that these mice are protected from hyperglycemia and progression to autoimmune diabetes. Importantly, depletion of CD122^+^ subset from NOD chimera derived CD45.2^+^CD8^+^ T cells before infusion led to peri-islet immune cell accumulation which further validates that the d-CD8^+^CD122^+^ subset from the NOD chimera are necessary to prevent autoreactive T cell attack of the islets of Langerhans.

### Donor-derived CD8^+^CD122^+^ Tregs generated in chimeric NOD mice have upregulated characteristic Treg markers

Given the superior protection conferred by d-CD8^+^CD122^+^ Tregs against diabetes progression, we performed a comprehensive phenotypic characterization to identify features associated with their protective capacity. The experimental design for d-CD8^+^CD122^+^ Treg phenotyping is depicted in **Fig 5a**. The d-CD8^+^CD122^+^ Tregs from chimeric mice were enriched for CD62L^−^CD44^+^ effector memory T (TEM) cells compared to CD8^+^CD122^+^ Tregs from WT C57BL/6 mice (43.75% versus 6.25%; p<0.0001) (**Fig. 5b-c**). The sustained enrichment of TEM-like d-CD8^+^CD122^+^ Tregs in the chimeric mice suggested an expanded, recall-ready Treg pool with capacity to rapidly exert regulatory suppression upon cognate antigen encounter, thereby promptly modulating effector response against the autoreactive T cells (22). Further flow cytometric analysis of d-CD8^+^CD122^+^ Tregs revealed upregulation of the transcription factor Helios, suggesting enhanced stability and suppressive functions of d-CD8^+^CD122^+^ Tregs compared to CD8^+^CD122^+^ Tregs from WT C57BL/6 mice (*p*<0.01). In addition, d-CD8^+^CD122^+^ Tregs showed upregulation of PD1, CD39, perforin, granzyme-B, ICOS, and FR4 compared to CD8^+^CD122^+^ Tregs from WT C57BL/6 mice (*p*<0.05). The d-CD8^+^CD122^+^ Tregs did not express the exhaustion marker TIM-3 (**Fig 5d)**. We further found that the bone marrow-derived d-CD8^+^CD122^+^ Tregs from the NOD chimeric mice exhibit a similar phenotype (**Fig S7**). Increased expression of these markers in d-CD8^+^CD122^+^ Tregs suggests that these Tregs have superior immune regulatory functions, including direct cytotoxicity promoting maintenance of immune tolerance in NOD chimera.

**Fig 5.**
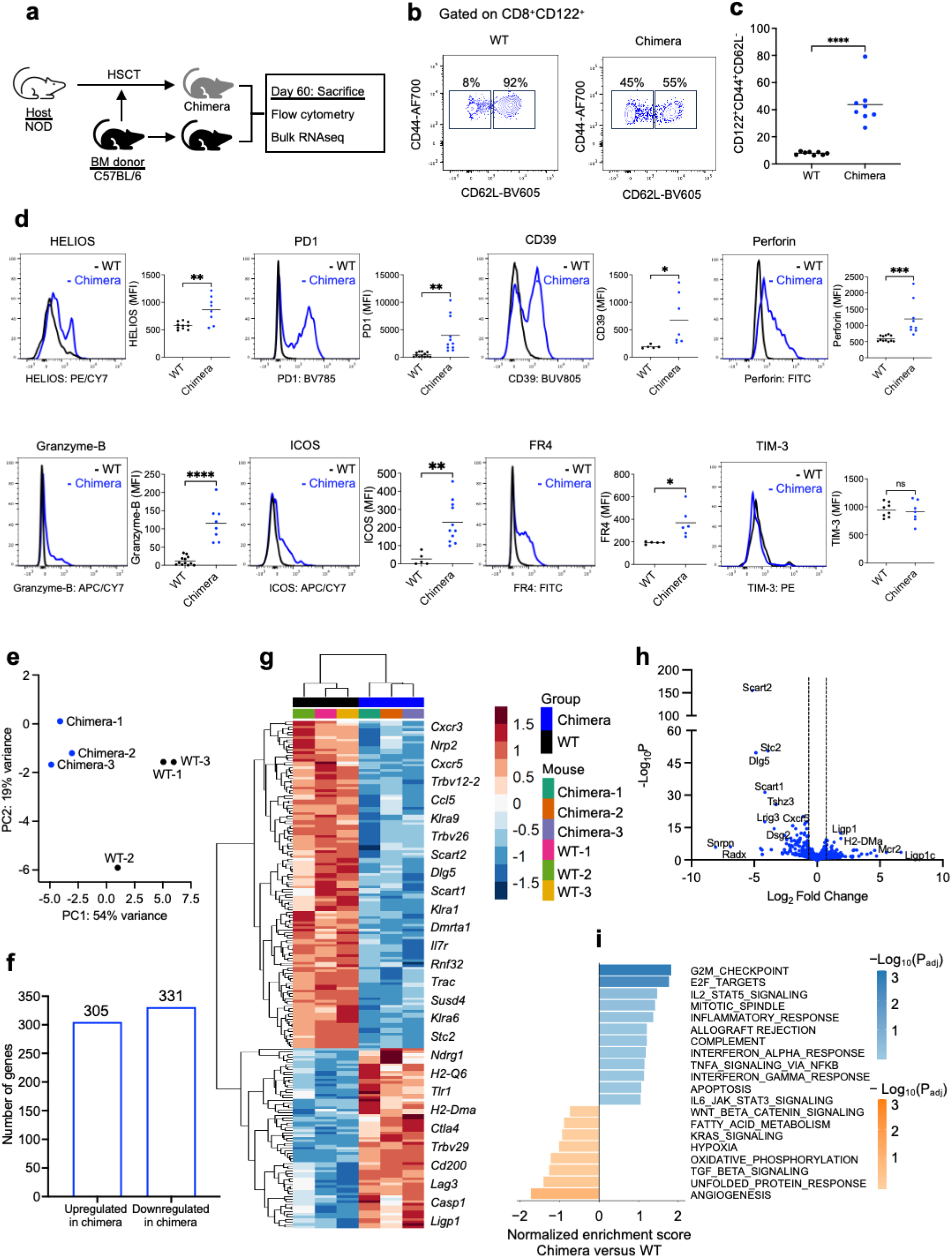
Donor-derived CD8^+^CD122^+^ Tregs have upregulated characteristic Treg markers. (**a**) Experiment design. NOD chimeras were sacrificed at day 60 post-HSCT and phenotype of CD45.2^+^CD122^+^ Tregs (d-CD8^+^CD122^+^ Tregs) was studied by flow cytometry analysis and bulk RNA sequencing. CD122^+^ Tregs from age and sex-matched WT C57BL/6 mice were used as controls. (**b**) Memory phenotype of d-CD8^+^CD122^+^ and WT CD8^+^CD122^+^ Tregs. (**c**) Quantification of percentage of CD44^+^CD62L^-^ effector memory from d-CD8^+^CD122^+^ and WT CD8^+^CD122^+^ Tregs. (**d**) Representative histograms showing expression of different suppressive markers in d-CD8^+^CD122^+^ and WT CD8^+^CD122^+^ Tregs. (**e**-**i**) Bulk RNA sequencing of d-CD8^+^CD122^+^ and WT CD8^+^CD122^+^ Tregs. (**e**) Principal component analysis of gene expression from d-CD8^+^CD122^+^ *versus* WT CD8^+^CD122^+^ Tregs. (**f**) Total number of upregulated and downregulated genes in d-CD8^+^CD122^+^ *versus* WT CD8^+^CD122^+^ Tregs with p_adj_ <0.1. (**g**) Heatmap showing differential gene expression in d-CD8^+^CD122^+^ *versus* WT CD8^+^CD122^+^ Tregs. (**h**) Volcano plot showing gene expression log_2_ fold change and –log_10_(P_adj_) in d-CD8^+^CD122^+^ *versus* WT CD8^+^CD122^+^ Tregs. Significant differentially expressed genes are defined by a log_2_ fold change > |1| and a P_adj_ < 0.001. (**i**) Gene-set enrichment analysis of d-CD8^+^CD122^+^ *versus* WT CD8^+^CD122^+^ Tregs. The 20 most immune-related and differentially expressed pathways by normalized enrichment score are shown. \*\*\*\**p*<0.0001. d-CD8^+^CD122^+^ Tregs; donor-derived CD8^+^CD122^+^ Tregs from NOD chimera, HSCT; hematopoietic stem cell transplantation, NOD; non-obese diabetic, WT; wildtype.

To further characterize d-CD8^+^CD122^+^ Tregs, we performed a detailed transcriptomic profiling of differences in gene expression of the d-CD8^+^CD122^+^ Tregs and WT C57BL/6 CD8^+^CD122^+^ Tregs. Principal component analysis revealed similarities among the samples within a group, while showed a significant variance in the gene profile between the CD8^+^CD122^+^ Tregs of chimeric NOD versus untreated donor C57BL/6 groups (**Fig 5e**). To further explore the differences in gene expression profiles in d-CD8^+^CD122^+^ Tregs, a differential gene analysis was performed, that identified a total of 305 upregulated genes and 331 downregulated genes (**Fig 5f**). Differential gene analysis of the top 200 genes demonstrated a notable increase in the expression of *Trbv29* and *Trbv12-2,* indicating a shift of TCR repertoire usage among d-CD8^+^CD122^+^ Tregs, suggesting an altered clonal representation. We found increase in the expression of *Ctla4*, *Cd200*, and *Lag3* in the d-CD8^+^CD122^+^ Tregs which indicated a more potent regulatory phenotype with stronger inhibitory signaling *via* CTLA4 and LAG-3 and enhanced suppression of T cell responses through CD200-CD200R-mediated inhibition. Notably, we found downregulation of Scavenger receptor family member expressed on T cells 1 and 2 (Scart1 and Scart2), which was further verified by qRT-PCR and flow cytometry (**Fig S8**). The downregulated expression of *Scart1 and Scart2*, which are typically associated with proinflammatory T cells (23), suggests a shift towards the generation of more resilient CD8^+^CD122^+^ Tregs in the chimeric mice. Upregulation of G2/M, and IL-2 signaling in the d-CD8^+^CD122^+^ Tregs suggested IL-2 dependent proliferation, while concurrent increases in inflammatory response, allograft rejection, IFN-α/IFN-γ, and TNF signaling *via* NF-κB reflects enhanced modulation of inflammatory cues (**Fig 5i**).

### Donor-derived CD8^+^CD122^+^ Tregs suppress autoreactive T cells *via* TCR CDR3 recognition

To elucidate the mechanism by which d-CD8^+^CD122^+^ Tregs control autoreactive T cells and confer protection against diabetes, we isolated IGRP-reactive CD8^+^ T cells from NOD mice and combined them with bulk NOD T cells to generate an IGRP-enriched T cell population. These cells were then cocultured for 72 h with CD8^+^CD122^+^ or CD8^+^CD122^−^ T cells isolated from either NOD mice or the donor-derived compartment of the chimeric mice. Representative flow cytometry plots showing the frequency of IGRP-reactive CD8^+^ T cells after coculture are presented in **Fig 6a**. Donor-derived CD8^+^CD122^+^ Tregs exhibited a greater capacity to reduce the frequency of IGRP-reactive CD8^+^ T cells than their NOD-derived CD8^+^CD122^+^ counterparts. In contrast, IGRP-reactive CD8^+^ T cells were not significantly affected by coculture with CD8^+^CD122^−^ T cells from either NOD or chimeric NOD mice (**Fig 6b**). Collectively, these findings indicate that d-CD8^+^CD122^+^ Tregs preferentially suppress the autoreactive CD8^+^ T cells, providing a mechanism for their protection against diabetes. Furthermore, in an *in vivo* T cell transfer experiment involving simultaneous transfer of WT NOD T cells and the Tregs into B6.129S-Rag2^tm1Fwa^ Cd47^tm1Fpl^ Il2rg^tm1Wjl^/J (triple KO C57BL/6) mice, we found that d-CD8^+^CD122^+^ preferentially suppress the IGRP-reactive CD8^+^ T cells (**Fig S9**). The use of triple KO mice in this experiment enabled us to dissect the autoreactive T cell-deletion function of d-CD8^+^CD122^+^ Tregs in the absence of alloreactive responses to host tissues.

**Fig 6.**
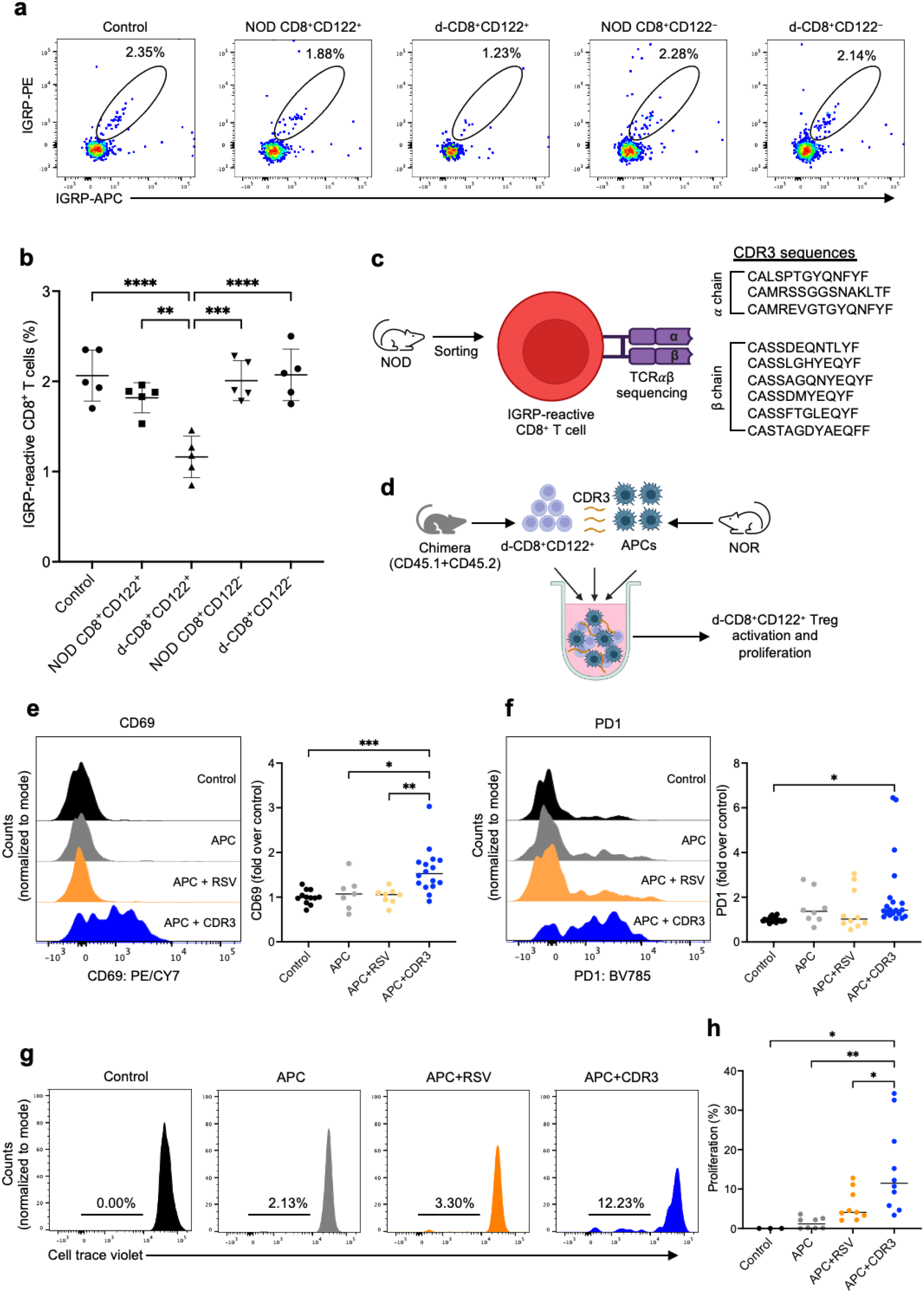
Donor-derived CD8^+^CD122^+^ Tregs suppress autoreactive T cells *via* CDR3 recognition. (**a**) Representative flow cytometry plots showing percentage of IGRP-reactive CD8^+^ T cells after coculture of IGRP-enriched NOD T cells with CD8^+^CD122^+^ Tregs or CD8^+^CD122^−^ T cells for 72h. (**b**) Quantification of percentage of IGRP-reactive CD8^+^ T cells after the coculture. (**c**) IGRP-reactive CD8^+^ T cells were sorted from NOD mice and T cell receptor αβ sequencing was performed. Multiple peptides from CDR3 sequences (3 from α chain and 6 from β chain) were synthesized. (RSV; SYIGSINNI) peptide sequence was used as an irrelevant antigen. (**d**) Design of Treg activation/proliferation assay. NOD chimeras were sacrificed at day 60 post-HSCT and CD45.2^+^CD8^+^CD122^+^ (d-CD8^+^CD122^+^) Tregs were sorted co-cultured with CD11b^+^ monocytes that were sorted from NOR mice in the presence of CDR3 peptide (sequence; CAMREVGTGYQNFYF). (**e-f**) Expression of surface markers CD69 and PD1 in d-CD8^+^CD122^+^ Tregs after 48 h of coculture. (**g**) Representative histograms showing dilution of CTV in d-CD8^+^CD122^+^ Tregs after 7 days of coculture. (**h**) Quantification of d-CD8^+^CD122^+^ Treg proliferation. Data were pooled from three independent experiments. \**p*<0.05, \*\**p*<0.01, \*\*\**p*<0.001, \*\*\*\**p*<0.0001. CDR3; complementarity determining region 3, CTV; cell trace violet, HSCT; hematopoietic stem cell transplantation, IGRP; islet-specific glucose-6-phosphatase catalytic subunit-related protein, NOD; non-obese diabetic, NOR; non-obese diabetes-resistant, RSV; respiratory syncytial virus.

These findings encouraged us to determine the molecular basis of IGRP-reactive CD8^+^ T cell killing by d-CD8^+^CD122^+^ Tregs. CD8^+^ T cells can recognize peptides presented by MHC-I molecules on the surface of target cells in order to maintain immune surveillance and to protect the body from intracellular pathogens (24). Endogenous self-derived peptide sequences can also be processed and presented on MHC-I molecules, enabling them to function as neoantigens for immune recognition (25, 26). CD8^+^ T cells have been shown to be able to control autoreactive T cells through the recognition of unique peptides generated from the TCR recombination area CDR3 of autoreactive T cells (27, 28). We hypothesized that d-CD8^+^CD122^+^ Tregs recognize CDR3-peptides derived from IGRP-reactive CD8^+^ T cells presented on MHC-I by APCs and/or IGRP-reactive CD8^+^ T cells themselves. This mechanism would enable selective and targeted clearance of IGRP-reactive CD8^+^ T cells to restore tolerance. To identify the unique TCR CDR3 peptides from IGRP-reactive CD8^+^ T cells targeted by d-CD8^+^CD122^+^ Tregs, we performed bulk TCR sequencing of IGRP-reactive CD8^+^ T cells and generated a series of corresponding CDR3 peptides derived from both *α* and β chain of TCR. We then performed *in vitro* co-culture involving d-CD8^+^CD122^+^ Tregs and NOR CD11b^+^ monocytes in the presence of 20 µM of CDR3 peptides for 48 h. We used NOR CD11b^+^ monocytes to enable presentation of the peptides in the identical MHC context as IGRP-reactive CD8^+^ T cells, allowing a direct, MHC-matched assessment of Treg interactions. We used 20 µM of RSV peptide (sequence: SYIGSINNI) as an irrelevant control (**Fig 6c-d, S10**). The introduction of a IGRP TCR α CDR3-derived peptide (sequence: CAMREVGTGYQNFYF) to the co-culture assay resulted in antigen-specific activation of d-CD8^+^CD122^+^ Tregs, as shown by an increase in the expression of activation markers CD69. In contrast, the introduction of RSV peptide to the co-culture resulted in a minimal non-specific background activation (**Fig 6e-f**). This data demonstrated that d-CD8^+^CD122^+^ Tregs specifically recognize and target IGRP-derived CDR3 peptides. Increased expression of co-inhibitory receptor PD1 in d-CD8^+^CD122^+^ Tregs was observed in the presence of CDR3 peptide, suggesting that the PD1-PDL1/PDL2 pathway may be utilized by the d-CD8^+^CD122^+^ Tregs in controlling autoimmune responses of IGRP-reactive CD8^+^ T cells. The activation of d-CD8^+^CD122^+^ Tregs was inhibited with the addition of MHC-I blocking antibodies (Pan MHC-I, H-2K^d^, or H-2D^b^) in the coculture system, but not with Qa-1 or MHC-II blocking antibodies (**Fig S11**). Furthermore, d-CD8^+^CD122^+^ Tregs proliferated as indicated by CTV dilution in a 7-day coculture system consisting of NOR CD11b^+^ monocytes and the CDR3 peptide (**Fig 6g-h, S12**). These data suggest that TCR CDR3 peptides of IGRP-reactive CD8^+^ T cells are presented on MHC-I of the residual IGRP-reactive CD8^+^ T cells, resulting in selective recognition and targeting of the IGRP-reactive CD8^+^ T cells by d-CD8^+^CD122^+^ Tregs. These data support fratricide of IGRP-reactive CD8^+^ T cells by d-CD8^+^CD122^+^ Tregs as mechanism of immunoregulation of autoreactive T cells in our mixed chimerism adoptive transfer models.

### CD8^+^CD122^+^ Tregs are impaired in individuals living with T1D

It is well-established that classical CD4^+^CD25^+^CD127^low^ Tregs from individuals with T1D are impaired in their suppressive function (7). These functional defects contribute to the loss of peripheral tolerance and T1D etiology. To better characterize the role of CD8^+^CD122^+^ Tregs in autoimmune diabetes, we undertook phenotypic studies of CD8^+^CD122^+^ Tregs from individuals with T1D to determine if they exhibit phenotypic or functional defects (**Fig 7a**). Live CD3^+^CD8^+^ T cells were identified by flow cytometric analysis of peripheral blood mononuclear cells (PBMCs) from individuals with T1D and healthy donors (HD). CD3^+^CD8^+^ T cells were further gated on CD122^+^ cells to determine the frequency of CD8^+^CD122^+^ Tregs (**Fig 7b**). We observed a statistically significant reduction in the frequency of CD8^+^CD122^+^ Tregs in T1D subjects compared to HD controls (0.74% versus 2.30%; *p*<0.01) (**Fig 7c**). The expression of Helios, a transcription factor required for Treg stability and function, was significantly reduced among CD8^+^CD122^+^ Tregs from T1D subjects (*p*<0.05) (**Fig 7d**). Our data are consistent with a previous study in T1D, wherein children with T1D were shown to have a significant reduction in the frequency of CXCR3^+^CD8^+^CD122^+^ Tregs (29). However, we did not observe any statistically significant differences in the expression of other surface markers tested including PD1, ICOS, CD38, HLA-DM, CD82, CD31, and CD200 (**Fig S13**). To evaluate the functionality of CD8^+^CD122^+^ Tregs, we performed *in vitro* suppression assay. We sorted CD3^+^ T cells and CD8^+^CD122^+^ Tregs from healthy donors and T1D patients. CD3^+^ T cells and Tregs were labeled with cell trace oregon green (CTOG) and cell trace violet (CTV), respectively, and co-cultured for 5 days to assess the suppressive function of the Tregs. The suppression assay demonstrated that CD8^+^CD122^+^ Tregs from patients with T1D exhibited impaired suppressive function compared with CD8^+^CD122^+^ Tregs from healthy donors (*p*<0.05 at 1:0.5 T cell to Treg ratio) (**Fig 7e**). These data suggest that reduced frequency and function of CD8^+^CD122^+^ Tregs may contribute to the pathology leading to T1D in humans.

**Fig 7.**
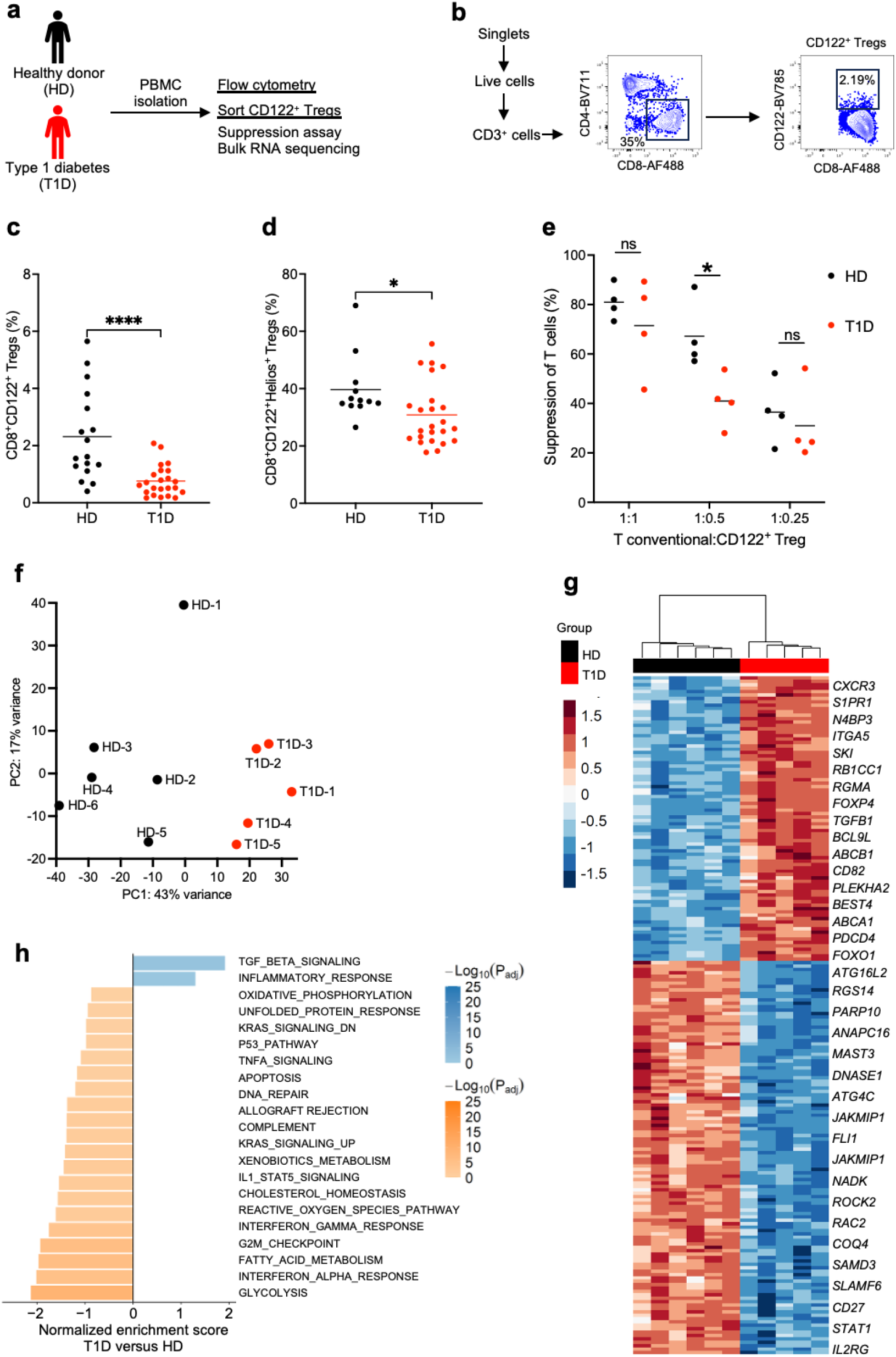
CD8^+^CD122^+^ Tregs are functionally impaired in individuals with T1D and have an altered transcriptomic signature. (**a**) Design of experiment. PBMCs from healthy donors (HD) and type a diabetes (T1D) patients were collected and the percentage of CD8^+^CD122^+^ among the CD8^+^ T cells and Helios^+^ within the CD8^+^CD122^+^ Tregs were analyzed by flow cytometry. For suppression assay and bulk RNA sequencing, CD8^+^ T cells were enriched, and CD8^+^CD122^+^ Tregs were sorted. (**b**) Gating hierarchy for human CD8^+^CD122^+^ Treg sorting. (**c**) Frequency of CD8^+^CD122^+^ Tregs among CD8^+^ T cells in HD and T1D. (**d**) Percentage of Helios-expressing cells among the CD8^+^CD122^+^ Tregs in HD and T1D. (**e**) Percent suppression of T conventional cells at different ratios of T conventional cells and CD8^+^CD122^+^ Tregs. (**f-h**) Bulk RNA sequencing of CD8^+^CD122^+^ Tregs from HD and T1D. (**f**) Principal component analysis of gene expression of CD8^+^CD122^+^ Tregs from HD and T1D. (**g**) Heatmap and hierarchical clustering of top 200 most differentially expressed genes in CD8^+^CD122^+^ Tregs by adjusted *p*-value in T1D *versus* HD. (**h**) Gene-set enrichment analysis of CD8^+^CD122^+^ Tregs from T1D *versus* HD. The 20 most immune-related and differentially expressed pathways by normalized enrichment score are shown. \**p*<0.05, \*\**p*<0.01. CD122^+^ Tregs; CD8^+^CD122^+^ Tregs, HD; healthy donors, PBMCs; peripheral blood mononuclear cells, T1D; type 1 diabetes.

To further characterize CD8^+^CD122^+^ Tregs in human T1D PBMC samples, we sorted CD8^+^CD122^+^ Tregs from individuals with T1D and HD and performed bulk RNA sequencing (**Table S1**). Principal component analysis revealed similarities among the samples within T1D subjects or HD while showed a significant variance in the gene profile between the CD8^+^CD122^+^ Tregs of T1D and healthy donors (**Fig 7f**). A differential gene expression analysis of CD8^+^CD122^+^ Tregs identified a total of 3648 upregulated and 4755 downregulated genes (**Fig S14**). Differential gene analysis of the top 200 genes showed significant downregulation of autophagy-related genes, notably *ATG16L2 and ATG4C.* This pattern indicates impaired autophagic flux, reducing the recycling of damaged organelles and proteins and thereby compromising cellular homeostasis in T1D-associated CD8^+^CD122^+^ Tregs. Moreover, the downregulation of *SLAMF7* and *CD27* signaling in CD8^+^CD122^+^ Tregs diminishes survival cues and Treg-T cell interactions, weakening suppressive capacity and stability in T1D. In addition, the concurrent downregulation of STAT1 and IL2RG in T1D-associated CD8^+^CD122^+^ Tregs suggests attenuated STAT1-dependent IFN-α/IFN-γ signaling and reduced γc cytokine signaling through IL2RG that drives Treg survival and function. We observed upregulation of genes such as *FOXP4, TGFB1, BCL9L, CD82, PDCD4, CD183,* and *IL21*, indicating a compensatory regulatory response to ongoing islet autoimmunity or a dysregulated or functionally impaired CD8^+^CD122^+^ Treg phenotype in T1D, potentially contributing to insufficient suppression or altered interactions with other immune cells such as autoreactive T cells (**Fig 7g-h**). Taken together, the functional and transcriptomic validation data indicate that CD8^+^CD122^+^ Tregs exhibit functional defects in T1D.

## DISCUSSION

Peripheral tolerance mechanisms are critical for restraining autoreactive cells that escape thymic negative selection. The potency of peripheral tolerance pathways in modulating disease is exemplified by the finding that many individuals exhibit islet autoantibodies and autoreactive T cells in peripheral blood without progressing to clinical T1D (30, 31). This observation suggests that additional protective mechanisms exist to maintain immune homeostasis and need to be better elucidated. There are a variety of cell populations that have been identified to mediate peripheral tolerance, including CD4^+^CD25^+^FOXP3^+^, CD8^+^CD158^+^, and type 1 regulatory (Tr1) cells (14, 32–35). One relatively recently identified CD8 population co-expresses CD122 and has a regulatory role in multiple settings such as islet allograft transplantation (11), contributing to the maintenance of immune tolerance and homeostasis (34, 36). CD8^+^CD158^+^ Tregs appear to uniquely regulate autoimmunity through the direct antigen-specific recognition and killing of autoreactive T cells (14, 37). In humans, CD8^+^CD158^+^ and CD8^+^CD122^+^ Tregs exhibit overlapping immunophenotypes, suggesting related regulatory lineages (**Fig S15**). In this study, we provide additional evidence that CD8^+^CD122^+^ Tregs play an important role in immune tolerance in T1D, as they are reduced and less functional in both NOD mice and humans with T1D. We also showed that induction of hematopoietic mixed chimerism revitalizes CD8^+^CD122^+^ Tregs in NOD recipients and directly prevents insulitis and hyperglycemia by targeted clearance of islet antigen-reactive CD8^+^ T cells *via* their TCR CDR3 recognition.

Our results revealed that CD8^+^CD122^+^ Tregs play a crucial role in abrogating autoreactivity in T1D. We showed generation of more suppressive donor-derived CD8^+^CD122^+^ Tregs in mixed chimeric NOD mice. These Tregs were phenotypically unique and functionally more competent while compared to their WT counterparts. Donor-derived CD8^+^CD122^+^ Tregs can differ in NOD chimera because they develop and persist in a distinct host environment, in which thymic selection, antigen exposure, and regulatory cues are altered compared to CD8^+^CD122^+^ Tregs from non-chimeric donor controls. In established chimeras, donor BM-derived Tregs may also acquire functional tolerance and adopt phenotypes that differ from donor controls, in part because they are shaped by recipient-derived antigen presenting cells. This divergence is likely further influenced by the conditioning regimen used to establish the chimera. For example, the total lymphoid irradiation conditioning used in this study could affect the selection, activation threshold, and functional state of d-CD8^+^CD122^+^ Tregs. In our study, the observed responsiveness of d-CD8^+^CD122^+^ Tregs to the CDR3 peptide suggest that the chimeric environment may enrich for peptide-reactive donor-derived clones without implying that the donor control repertoire is intrinsically identical in phenotype or specificity. Comprehensive phenotyping of the d-CD8^+^CD122^+^ Tregs revealed an upregulation of key immune-suppressive markers, including Helios, PD1, and ICOS. It has been established that Helios-dependent STAT5 activation is required for CD8^+^ Treg survival and function (38). Furthermore, the results showing reduced expression of scavenger receptors Scart1 and Scart2 in d-CD8^+^CD122^+^ Tregs may serve as informative markers to distinguish CD8^+^CD122^+^ Tregs. Taken together, CD8^+^CD122^+^Helios^+^PD1^+^Scart1/2^-^ Tregs represent a novel and distinct Treg lineage involved in maintaining peripheral tolerance to autoimmunity. Further functional validation including suppression assays, Treg stability assessments, and *in vivo* tolerance models-is required to better characterize the role of these Tregs in autoimmunity and transplantation settings.

A previous study reported impaired suppressive function of CD8^+^CD122^+^ Tregs in NOD mice (8). In this study, we confirmed both loss of function and reduced numbers of CD8^+^CD122^+^ Tregs. this could be possibly due to multiple mechanisms including defective thymic development, inflammatory milieu, and competition of growth signals (e.g. IL-2) in the autoimmune environment. Our human findings are also consistent with a previous study that found a significant reduction in the frequency of CXCR3^+^CD8^+^CD122^+^ Tregs in children with T1D (29). Our results contrast with other autoimmune diseases, including systemic lupus erythematosus, multiple sclerosis, and celiac diseases, wherein the CD8^+^CD158^+^ Tregs were found to be increased (14). In addition, a recent study shows an increased frequency of CD8^+^CD158^+^ Tregs during pregnancy and suggests that this subset contributes to maternal-fetal tolerance by maintaining fetal antigen-specific alloreactive T cell responses (39). This discrepancy is likely due to disease-specific genetic, immunological, and environmental conditions that distinguishes T1D etiology and the use of disease-modifying medications to treat other autoimmune conditions. The T1D subjects in our study included individuals with established T1D receiving exogenous insulin therapy and are not receiving other immune modifying therapies. It is possible that evaluation of CD8^+^CD122^+^ may help inform progression from stage 1 or 2 to later disease stage.

In the context of CD4^+^ Tregs, professional antigen presenting cells are required to present peptides on MHC-II molecules and the recognition of these MHC-II bound peptides occurs through the TCR of CD4^+^ Tregs (40). In contrast, CD8^+^ T cells can recognize peptides presented on MHC-I, which is expressed on nearly all nucleated cells (41). This recognition triggers the release of cytotoxic molecules (i.e. perforin, granzyme-B) from CD8^+^ T cells to directly kill target cells or to recruit other immune cells to the site of inflammation (42, 43). We observed that d-CD8^+^CD122^+^ Tregs were activated upon coculture with antigen presenting cells in the presence of peptides derived from the CDR3 region of TCR α chain of autoreactive T cells. Previous studies have suggested that endogenous T cell peptides can be presented on MHC-I molecules, allowing for unique recognition by CD8^+^ T cells (41, 44) and that the CD8^+^ T cells from T1D subjects fail to recognize peptide expressed on target cells, suggesting that this defect may play a major in the loss of peripheral tolerance in T1D (41). In the present study, we have established that d-CD8^+^CD122^+^ Tregs possess the ability to specifically suppress autoreactive IGRP-reactive CD8^+^ T cells through the recognition of peptides derived from unique CDR3 regions. MHC-I molecules typically present 8-10 amino acids in length (45). These peptide-MHC I complexes can be directly recognized by the T cell receptor of CD8^+^ T cells, with the CD8 coreceptor stabilizing the interaction and facilitating downstream signaling. This unique mechanism of target cell recognition is a distinguishing factor for CD8^+^ Tregs since they would have very high specificity in their control of autoimmunity and would not be globally suppressive, which is potentially beneficial for precise immunotherapies. In our study, although the CDR3-derived peptide used to activate the Tregs is considerably larger for MHC-I loading, it is plausible that the 15-mer peptide may undergo intracellular processing following uptake by antigen presenting cells, generating shorter peptide fragments that can be presented by MHC class I molecules and recognized by d-CD8^+^CD122^+^ Tregs. Our findings further suggest that the d-CD8^+^CD122^+^ Tregs generated in chimeric NOD mice constitute a polyclonal population, with distinct clones potentially recognizing one or more CDR3 derived epitopes from autoreactive T cells. In addition, the antigen-specific activation of a subset of these Tregs may create a cytokine- and costimulation-rich environment that promotes the bystander activation or expansion of other d-CD8^+^CD122^+^ Tregs. Thus, the expansion observed *in vitro* may reflect the combined contributions of antigen-specific recognition and bystander amplification rather than the selective expansion of a population with a single antigen specificity.

The limitations of this study should be acknowledged. First, our analyses were performed using the total CD8^+^CD122^+^ Treg population, which likely represents a heterogenous compartment comprising both regulatory and non-regulatory cells rather than a discrete Treg lineage. Additional markers such as HELIOS, PD1, CD39, Ly49, and CD49d, may help further dissect the specific population responsible for the observed regulatory effects. This will require subset-specific functional studies in future. Second, although CD8^+^CD122^+^ Tregs responded to CDR3-derived 15-mer peptide sequence, this peptide is likely processed intracellularly into shorter MHC class I-restricted epitopes prior to presentation. Thus, the precise antigenic determinants recognized by CD8^+^CD122^+^ Tregs remain to be identified. Future studies combining MHC class I-bound peptide elution, epitope mapping, and peptide-MHC tetramer approaches will define the naturally presented CDR3 epitopes, enable isolation of antigen-specific CD8^+^CD122^+^ Treg subsets, and facilitate their phenotypic and functional characterization. Finally, while our findings establish a mechanistic framework in experimental models, validation of CD8^+^CD122^+^ Tregs and their antigen-specific subsets in human samples will be essential to establish their translational therapeutic potential in autoimmune diabetes.

In conclusion, our study highlights a promising strategy to restore the loss of CD8^+^CD122^+^ Treg functionality and a molecular mechanism of autoreactive T cell recognition by d-CD8^+^CD122^+^ Tregs which are generated in mixed chimerism. Our findings suggest a therapeutic strategy to restore CD8^+^CD122^+^ Tregs that precisely target autoreactive T cells *via* TCR CDR3 recognition. The decreased frequency and functional impairments observed in these Tregs in both murine and human contexts underscore the need for further research into targeted therapies that could restore the Treg function and maintain peripheral tolerance in T1D. Investigation of potential strategies to restore the functional deficits in CD8^+^CD122^+^ Tregs opens an avenue towards revitalizing the Tregs to restore their immunoregulatory functions. We hypothesize that simultaneous loss of the CD4^+^CD25^+^ and CD8^+^CD122^+^ Treg subsets contributes to the loss of peripheral tolerance and accelerates autoimmunity in T1D. Additionally, it could be postulated that other Treg subsets, such as type 1 regulatory T cells and other less well characterized subtypes, may also be dysregulated in autoimmune diseases including T1D and warrant investigation in future studies.

## MATERIALS AND METHODS

### Animals

All experimental mice were purchased from Jackson Laboratory (Sacramento, CA) and housed in Stanford University Animal Facility. Animal experiments were performed according to the protocols approved by Institutional Animal Care and Use Committee of Stanford University. Prediabetic female NOD/ShiLtJ mice (stock no. 001976) were used as BM recipients and C57BL/6J (stock no. 000664) were used as BM donors. For adoptive T cell transfer experiments, NOD-Rag1^null^IL2rg^null^ (NRG; stock no. 007799) and B6.129S-Rag2^tm1Fwa^Cd47^tm1Fpl^ Il2rg^tm1Wjl^/J (Triple KO; stock no. 025730) mice were used. NOR/LtJ (stock no. 002050) were used as controls for some experiments.

### Generation of mixed donor chimerism in NOD mice

NOD mice were preconditioned with a non-myeloablative conditioning regimen developed by our group (17). Briefly, prediabetic female NOD mice (10 weeks old) were preconditioned with 8 doses of total lymphoid irradiation (TLI: 2.4 Gy), 2 doses of total body irradiation (TBI: 1.5 Gy) using a Kimtron Biological Irradiator (Oxford, CT), and 5 doses of anti-thymocyte serum (ATS: 50 μL, Accurate Chemical Supplies, Carle Place, NY). Femoral and tibial bones were collected from donor C57BL/6 mice and bone marrow (BM) suspensions were prepared by flushing the bones with RPMI-1640 media containing 10% fetal bovine serum. Cell suspensions were filtered through sterile 70 *µ*m nylon mesh strainers, washed, and resuspended in sterile phosphate-buffered saline. A total of 50×10^6^ whole BM cells were intravenously injected into the preconditioned NOD mice *via* retroorbital injection. Donor chimerism was then analyzed at day 60 post HSCT using method reported previously (17).

### Adoptive T cell transfer and blood glucose monitoring

The NOD chimeras were euthanized at day 60 post-BMT and CD3^+^ T cells were sorted to purity using a FACSAria II at the Stanford Shared FACS facility. Then, 2 × 10^6^ bulk or CD25-depleted CD3^+^ T cells were intravenously injected into NRG mice. As a positive control for diabetes induction, some NRG mice were injected with 2 × 10^6^ CD3^+^ T cells sorted from 12-14 weeks old prediabetic wildtype (WT) NOD mice. To demonstrate the protective role of d-CD8^+^CD122^+^ Tregs, we simultaneously injected 3 × 10^6^ CD3^+^ T cells from 12-14 weeks old prediabetic WT NOD mice and 2 × 10^6^ bulk or CD122 depleted CD8^+^ T cells from NOD chimera or WT C57BL/6 mice. Then, the recipient NRG mice were monitored for diabetes incidence by weekly checking their non-fasting blood glucose levels using the Contour Next EZ blood glucose monitoring system (Ascensia Diabetes Care, Parsippany, NJ).

### Flow cytometry

Antibodies were purchased from BioLegend unless otherwise stated. Live cells from singlet gating were identified by Zombie Aqua™ (BioLegend) and stained with antibodies at a final dilution of 1:200. The following antibodies were used for chimerism analysis and murine CD122^+^ Treg characterization: Brilliant Violet 711 (BV711)-conjugated CD45.1 (clone A20), Pacific Blue (PB)-conjugated CD45.2 (clone 104), Brilliant Ultraviolet 737 (BUV737)-conjugated CD3 (BD Biosciences; clone 145-2C11), Allophycocyanin Cyanine 7 (APC/Cy7)-conjugated B220 (clone RA3-6B2), BUV805-conjugated CD8 (BD Biosciences; clone H35-17.2), BUV395-conjugated CD4 (ThermoFisher Scientific; clone RM4-5), BUV563-conjugated CD25 (ThermoFisher Scientific; clone PC61.5), Alexa Fluor 700 (AF700)-conjugated CD44 (clone IM7), Phycoerythrin (PE)-conjugated CD122 (clone 5H4), BV605-conjugated CD62L (clone MEL-14), BV785-conjugated PD1 (clone 29F.1A12), Fluorescein isothiocyanate (FITC)-conjugated perforin (clone S16009A), APC/Cy7 conjugated Granzyme B (clone QA16A02), PE/Cy7-conjugated Helios (clone 22F6), FITC-conjugated FR4 (clone 12A5), PE/CY7-conjugated CD69 (clone H1.2F3), APC-conjugated FASL (clone MFL3), Percp/Cy5.5-conjugated TIM-3 (clone B8.2C12), APC/CY7-conjugated ICOS (clone C398.4A), and PE/CY7-conjugated CD38 (clone 90).

The following antibodies were used for CD8^+^CD122^+^ Treg characterization from human samples: BUV805-conjugated CD3 (BD Biosciences; clone UCHT1), BV711-conjugated CD4 (clone OKT4), BUV661-conjugated CD25 (BD Biosciences; clone M-A251), RB670-conjugated Foxp3 (BD Biosciences; clone 259D/C7), APC/H7-conjugated CD8 (BD Biosciences; clone SK1), BV786-conjugated CD122 (BD Biosciences; clone Mik-B3), DyLight^TM^ 405-conjugated CD163L1 (Novus Biologicals; clone CD163L1/7972), BV605-conjugated CD158 (BD Biosciences; clone DX9), Peridinin Chlorophyll Protein Cyanine 5.5 (PerCP/Cy5.5)-conjugated Helios (clone 22F6), AF700-conjugated PD-1 (clone EH12.2H7), PE-conjugated HLA-DM (clone MaP.DM1), and PE/Cy7-conjugated CD39 (clone A1), PE/Dazzle-conjugated CD31 (clone WM59), BV421-conjugated CD200 (clone OX-104), APC-conjugated CD82 (clone ASL-24), and BV711-conjugated CD38 (clone HIT2). The following antibodies were used for sorting human CD122^+^ Tregs: PE-conjugated CD3 (clone OKT3), BV711-conjugated CD4 (clone OKT4), AF488-conjugated CD8 (clone SK1), BV786-conjugated CD122 (BD Biosciences; clone Mik-B3). Data were collected using BD FACSymphony Cell Analyzer and analyzed by FlowJo Software Version 10.9.0 (Tree Star, Ashland, OR).

### Suppression assay

Live CD3^+^ T cells (responders) and CD8^+^CD122^+^ Tregs (suppressors) were sorted to purity using BD FACS Aria II cell sorter. Then, the responder T cells were labeled using a CellTrace^TM^ Violet (CTV) Proliferation Kit (ThermoFisher) and cocultured with the suppressor CD8^+^CD122^+^ Tregs in the presence of Dynabeads^TM^ Mouse T-Activator CD3/CD28 beads (ThermoFisher) for 5 days in a 96-well U-bottom tissue culture plate (Corning, Glendale, AZ). The percentage of T cell suppression was calculated by measuring CTV dilution in the responder T cells by flow cytometry.

### Detection of autoreactive T cells

Islet-specific glucose-6-phosphatase subunit-related protein (IGRP)-reactive CD8^+^ CTL were identified by tetramer staining. APC and PE-conjugated IGRP H-2^kd^ tetramers (Sequence 206-214: VYLKTNVFL) were obtained from the NIH Tetramer Core Facility (Bethesda, MD) and used at a final concentration of 1:200 at 4°C for 30 min. The cells were washed, stained further for CD3 and CD8 surface markers, and analyzed by flow cytometry.

### Histological analysis

Hematoxylin and eosin staining, and immunohistochemistry of the pancreas was performed at day 60 post-HSCT. Mice were euthanized and pancreata were isolated and fixed in formalin before embedding in paraffin blocks. Sections were then stained with hematoxylin and eosin using method described previously (46). Immunohistochemistry staining was performed using methods reported previously (17, 46, 47). The following antibodies were used for immunohistochemistry analysis: anti-CD3 (Cell Signaling, clone D7A6E, dilution 1:200) and CD68 (Cell Signaling, clone E3O7V, dilution 1:150).

### Bulk RNA and TCR sequencing

Total RNA was isolated using a NucleoSpin RNA Kit (Ref: 740955.50) and quantified using a Qubit Fluorometer (Invitrogen) and TapeStation 4200 (Agilent Technologies) for quality assessment. RNA libraries were prepared using the Takara SMART-Seq mRNA Kit (Cat. #634773) following the manufacturer’s instructions. Libraries were constructed using the Nextera XT DNA Library Preparation Kit (Illumina, Cat. #FC-131-1096) and indexed *via* tagmentation and PCR amplification to generate Illumina-compatible libraries. Indexed libraries were pooled, normalized to 4 nM, and sequenced on an Illumina NovaSeq X Plus platform using paired-end (2×100 bp) reads. T-cell receptor (TCR) libraries were prepared using the Takara SMART-Seq® Mouse TCR (with UMIs) Kit (Cat. #634857) following the manufacturer’s instructions. The resulting full-length cDNA molecules were PCR-amplified to enrich for TCR sequences and purified using Agencourt AMPure XP beads (Beckman Coulter). Purified cDNA was then used to generate Illumina-compatible sequencing libraries through indexed PCR amplification. Library quality and quantity were confirmed using the TapeStation 4200 (Agilent Technologies) and Qubit Fluorometer (Invitrogen). Indexed libraries were normalized to 4 nM, pooled, and sequenced on an Illumina MiSeq platform using paired-end (2×300 bp) reads.

All sequencing analysis was performed in RStudio with R4.4. For RNAseq analysis, after loading data into DESeq2 (48), genes with fewer than 10 reads across all samples were removed. After standard quality control, differential expression analysis of human samples was performed using the formula∼ T1D status. Additional variables that were considered in the analysis included age, biological sex, ethnicity, and year of diagnosis. This information was not available for healthy donors, however, and therefore was excluded from the differential expression analysis. For murine samples, differential expression analysis was performed using the formula∼ strain or chimera status. Gene-set enrichment was performed in fgsea (49) with 100,000 permutations using the Hallmark gene sets (available at MSigDB) and selected statistically significant (padj<0.25), immunologically-related pathways were included for visualization. TCR sequencing analysis was performed using immunarch after downsampling to the smallest number of total clones within each experiment.

### Quantitative RT-PCR

CD8^+^CD122^+^ Tregs were sorted to purity and lysed in triazole (Thermofisher Scientific). RNA extraction was then done using an RNA mini kit (Qiagen, Germany) according to the manufacturer’s guidelines and the quality of RNA was evaluated using an Agilent bioanalyzer 2100 system. Paired-end 150 base pair sequencing was carried out using an Illumina NovaSeq platform. The quality assessment, read filtering, and mapping were performed using the NGS QC toolkit and alignment was performed using HISTAT2 tool against Mus Musculus (GRCm39/mm39). HTSeq was used to quantify the reads. For RT-PCR, iTaq Universal One-Step RT-qPCR Kits were used according to the manufacturer’s guidelines. The following primers were used: SCART1 (forward primer: TGGACTTCGTGCGGCCATGA-3’; backward primer: 5’-CTACTCGTCATTCTCTGTGG-3’), and SCART2 (forward primer: 5’-TGTGAACGTCCTDDAAGT-3’; backward primer: 5’-GGGCTTCCACGTGACCTGT-3’).

### Treg activation and proliferation assays

IGRP-reactive CD8^+^ CTL were sorted from prediabetic NOD mice and processed for bulk TCR sequencing. Multiple peptide sequences targeting the TCR CDR3 regions of both α and β chains were generated. The synthesis and purification of these peptides was conducted by ELIM Biopharmaceuticals (Hayward, CA). Sorted d-CD8^+^CD122^+^ Tregs (1 × 10^5^) from chimeric mice were cocultured with CD11b^+^ monocytes (2 × 10^5^) isolated from NOR mice in the presence of 20 µM CDR3 peptides for 48 h (15-mer length, sequence CAMREVGTGYQNFYF). Respiratory Syncytial Virus peptide sequence (RSV; SYIGSINNI) was used as an irrelevant antigen. For proliferation assays, d-CD8^+^CD122^+^ Tregs were sorted, CTV-labeled, and co-cultured with NOR CD11b^+^ monocytes. A total of 2 × 10^4^ d-CD8^+^CD122+ Tregs were co-cultured with 1 × 10^5^ CD11b^+^ monocytes in the presence of 20 of µM RSV or CDR3 peptide for 7 days. Dilution of CTV in d-CD8^+^CD122^+^ Tregs was analyzed by flow cytometry.

### Evaluation of CD8^+^CD122^+^ Tregs in individuals with T1D

Individuals with T1D provided informed consent in accordance with the Declaration of Helsinki and were enrolled in a research protocol approved by the Stanford University Institutional Review Board (IRB) protocol #35453. Blood samples were collected, and peripheral blood mononuclear cells (PBMCs) were isolated by density gradient centrifugation using Ficoll-Paque (Cytiva, Marlborough, MA). Healthy donor blood samples were obtained from the Stanford Blood Center (Palo Alto, CA) from donors aged 18-65 years; additional demographic information was not available to the investigators because samples were provided in a de-identified manner to protect donor privacy. Blood samples from individuals with T1D were obtained through the Stanford T1D clinic. T1D diagnosis was confirmed by the presence of two or more islet autoantibodies documented at the clinical evaluation. At the time of blood collection, all participants had been diagnosed with T1D for more than one year and were receiving insulin therapy.

### Statistical analysis

Statistical analysis was performed by using GraphPad Prism 10 (La Jolla, CA). Statistical values were calculated using unpaired *t*-test or one-way ANOVA. Differences with *p* values less than 0.05 were considered statistically significant.

## Supplementary Materials

**Fig S1.** Gating hierarchy of murine CD8^+^CD122^+^ Tregs.

**Fig S2.** Phenotypic characterization of CD8^+^CD122^+^ Tregs from NOR and NOD mice.

**Fig S3.** Details of the conditioning regimen used for HSCT.

**Fig S4.** Detection and characterization of IGRP-reactive CD8^+^ T cells.

**Fig S5.** Donor chimerism in peripheral blood, spleen, and pancreas.

**Fig S6.** Host CD25^+^ T cells isolated from NOD chimera prevent diabetes onset.

**Fig S7.** Phenotype of bone marrow-derived d-CD8^+^CD122^+^ Tregs.

**Fig S8.** Donor-derived CD8^+^CD122^+^ Tregs have reduced expression Scart1 and Scart2.

**Fig S9.** Donor-derived CD8^+^CD122^+^ Tregs preferentially deplete autoreactive T cells.

**Fig S10.** Screening of CDR3 peptides using activation assay.

**Fig S11.** MHC-I blockade abrogated CDR3 peptide-mediated activation of d-CD8^+^CD122^+^ Tregs.

**Fig S12.** Proposed mechanism of IGRP-reactive CD8^+^ T cell killing by d-CD8^+^CD122^+^ Tregs *in vitro*.

**Fig S13.** Phenotype of CD8^+^CD122^+^ Tregs in individuals with T1D.

**Fig S14.** Bulk RNA sequencing of human CD8^+^CD122^+^ Tregs.

**Fig S15.** Representative flow cytometry plot showing overlapping phenotypic characteristics between CD8^+^CD122^+^ Tregs and CD8^+^CD158^+^ Tregs in human sample.

## ABBREVIATIONS

ATS: Anti-thymocyte serum
APC: Antigen-presenting cell
BM: Bone marrow
CD: Cluster of differentiation
CDR3: Complementarity-determining region-3
CTV: Cell trace violet
FR4: Folate receptor 4
HSCT: Hematopoietic stem cell transplantation
ICOS: Inducible T-cell costimulator
IFN-α: Interferon alpha
IFN-γ: Interferon gamma
IGRP: Islet-specific glucose-6-phosphatase subunit-related protein
IL-2: Interleukin 2
KIR: Killer-like immunoglobulin-like receptor
MHC: Major histocompatibility complex
NOD: Non-obese diabetic
NOR: Non-obese diabetes-resistant
NRG: NOD-Rag1^null^IL2rg^null^
PD1: Programmed cell death protein 1
SCART: Scavenger receptor family member expressed on T cells
SSC-A: Side scatter area
T1D: Type 1 diabetes
TBI: Total body irradiation
TCM: T central memory
TCR: T cell receptor
TEM: T effector memory
TIM-3: T-cell immunoglobulin and mucin-domain containing-3
TLI: Total lymphoid irradiation
Treg: Regulatory T cell
WT: wildtype

## Acknowledgements

We thank the staff of the Stanford University Animal Care Facility for taking exceptional care of the mice during the experiments. BD FACSymphony flow cytometer was purchased with funds from an NIH Shared Instrumentation (grant no. 1S10OD026831-01).

## Author contribution

Conceptualization: SP, EHM

Methodology: SP, CSB, SR, KPJ, MMD, EHM

Investigation: SP, CSB, BPI, SR, PC, BG, XW, BM, AW, LW, KJ, WH, ES, AT, RA, SD, NN

Visualization: SP, CSB, SR, KPJ, MMD, EHM

Feedback: GF, AST, KPJ, MMD, EHM

Funding acquisition: AST, EHM

Project administration: KPJ, EHM

Supervision: EHM

Writing-original draft: SP

Writing-review and editing: SP, KPJ, MMD, EHM

## Data and materials availability

The raw and processed sequencing data are available in the GEO repository under accession number GSE338315. All other data are available in the main text or the supplementary materials.

