## Supplementary Information for "Donor-derived CD8^+^CD122^+^ Tregs generated in mixed donor chimeric NOD mice suppress autoreactive T cells"

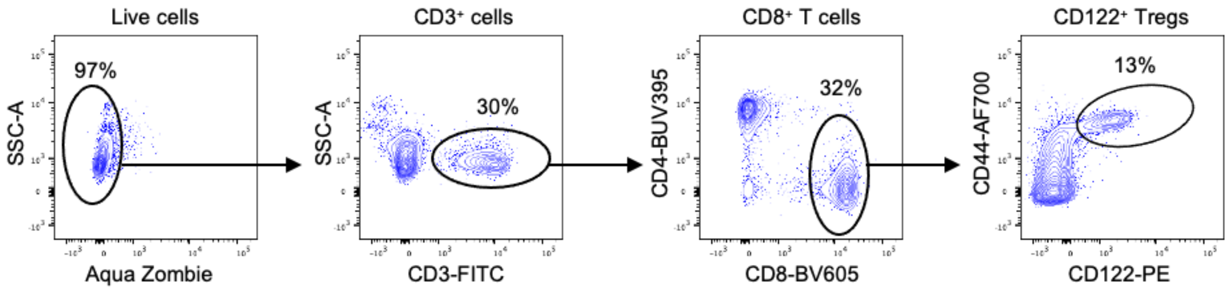


**Fig S1.** Gating hierarchy of murine CD8$^+$CD122$^+$ Tregs. Dead cells were excluded by staining with Zombie Aqua™ Fixable Viability Kit and CD3$^+$CD8$^+$CD44$^+$CD122$^+$ Tregs were gated from the live cells.

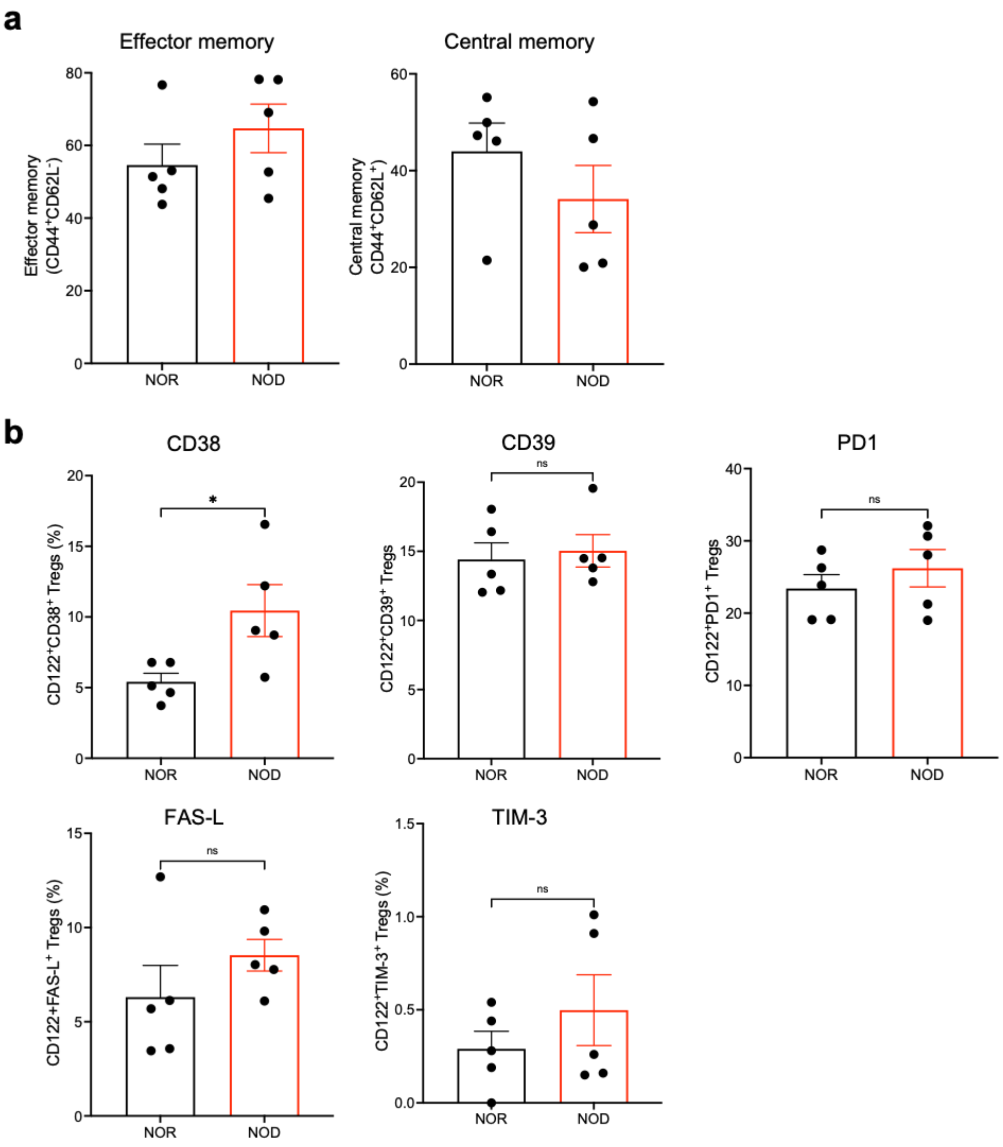


**Fig S2.** Phenotypic characterization of CD8$^+$CD122$^+$ Tregs from NOR and NOD mice. (**a**) Percentage of effector memory (CD44$^+$CD62L$^-$) Tregs and central memory (CD44$^+$CD62L$^+$) CD8$^+$CD122$^+$ Tregs in NOR and NOD. (**b**) Expression of different surface markers in NOR and NOD CD8$^+$CD122$^+$ Tregs. **p*<0.05. ns: not significant. FAS-L: Fas ligand, NOD: non-obese diabetic, NOR: non-obese diabetes-resistant, PD1: programmed cell-death protein 1, TIM-3: T-cell immunoglobulin and mucin-domain containing-3.

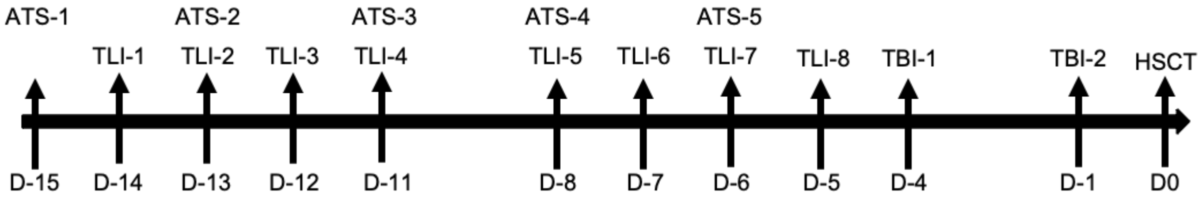


**Fig S3.** Details of the conditioning regimen used for HSCT. Prediabetic NOD mice were conditioned with 8 doses of total lymphoid irradiation (2.4 Gy each dose), two doses of total body irradiation (1.5 Gy each dose), and 5 doses of anti-thymocyte serum (50 $\mu$L each dose). TLI was given at day -14, -13, -12, -11, -8, -7, -6, and -5 prior to HSCT. TBI was given at day -4 and day -1 prior to HSCT. ATS was given at day -15, -13, -11, -8, and -7 prior to HSCT. A total of $50 \times 10^6$ whole bone marrow cells from C57BL/6 was injected into each mouse at day 0. ATS: antithymocyte serum, D: day, HSCT: hematopoietic stem cell transplantation, NOD: non-obese diabetic, TBI: total body irradiation, TLI: total lymphoid irradiation.

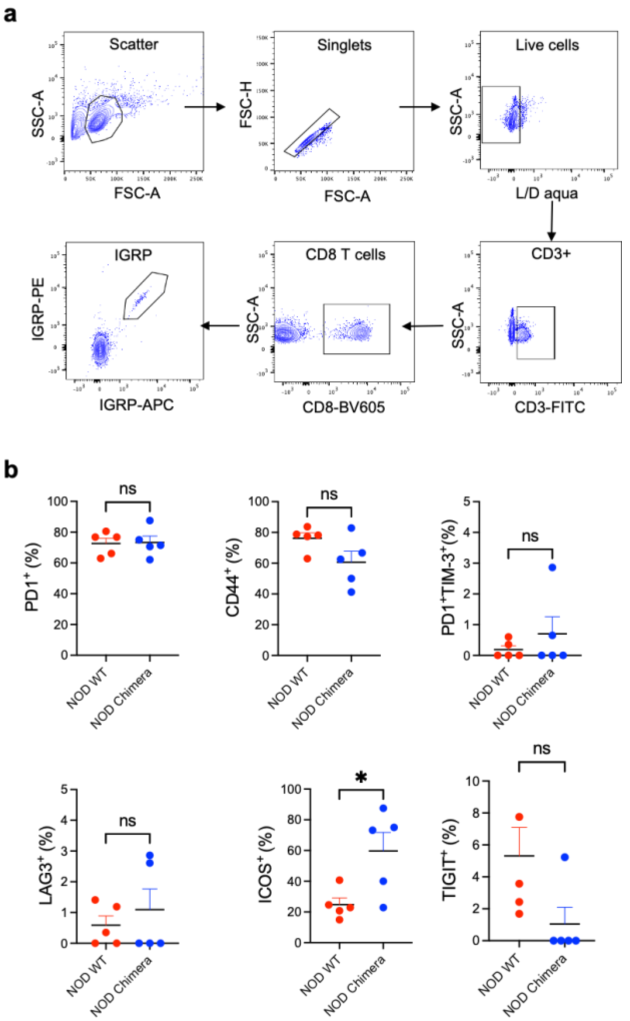


**Fig S4**. Detection and characterization of IGRP-reactive CD8$^+$ T cells. (**a**) Gating hierarchy of IGRP-reactive CD8$^+$ T cells in NOD mice. Dead cells were excluded by staining with Zombie Aqua™ Fixable Viability Kit. CD3$^+$CD8$^+$ T cells that were double-positive for IGRP-APC and IGRP-PE tetramers were considered IGRP-reactive CD8$^+$ T cells. (**b**) Expression of different surface markers in IGRP-reactive CD8$^+$ T cells in NOD WT and NOD chimera. *$p$<0.05. ns: not significant. ICOS: inducible T-cell costimulator, IGRP: islet-specific glucose-6-phosphatase subunit-related protein, LAG3: lymphocyte activation gene 3, PD1: programmed cell-death protein 1, TIGIT: T cell immunoreceptor with Ig and ITIM domains, TIM-3: T-cell immunoglobulin and mucin-domain containing-3, WT: wildtype.

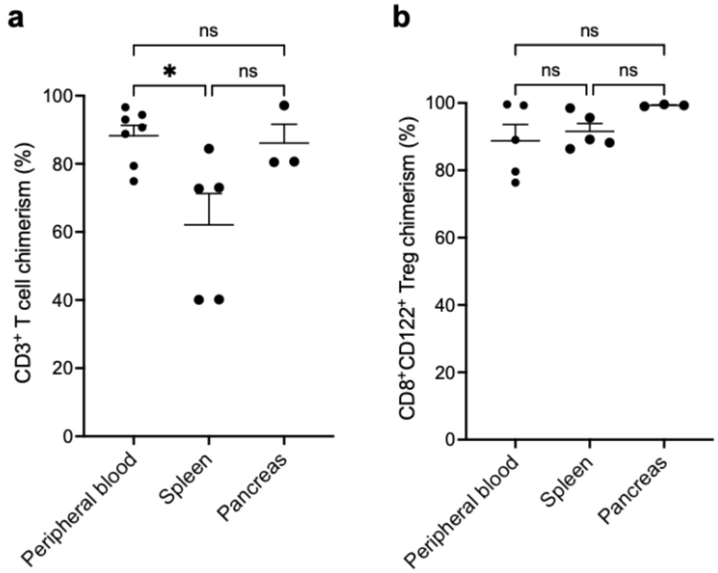


**Fig S5.** Analysis of donor chimerism in peripheral blood, spleen, and pancreas at day 60 post-HSCT. To analyze chimerism in pancreas, NOD chimeras were euthanized at day 60 post-HSCT, pancreata were collected and digested using 1 mg/mL collagenase IV (Millipore Sigma) at 37°C for 10 min. Lymphocytes were isolated from digested pancreata by density gradient centrifugation using Ficoll-Paque™ PLUS (Cytiva). Isolated lymphocytes were stained with antibody against CD3, CD8, CD44, CD122, CD45.1, and CD45.2. Flow cytometry was used to analyze T cells in pancreata. (a) $CD3^+$ T cell chimerism. (b) $CD8^+CD122^+$ Treg chimerism. ns: not significant, $*p<0.05$.

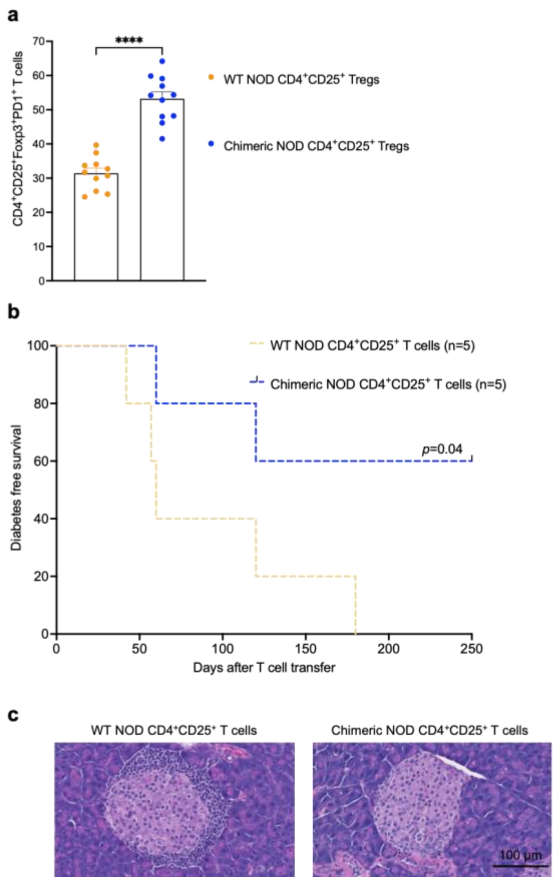


**Fig S6.** (**a**) Elevated PD1 expression in host-derived CD4$^+$CD25$^+$ Tregs of chimeric mice. (**b-c**) Adoptive T cell transfer and monitoring of diabetes induction in NRG mice. Host CD25$^+$ T cells isolated from NOD chimera prevent diabetes onset. NOD T cells ($2 \times 10^6$ per mouse) and CD4$^+$CD25$^+$ T cells from WT NOD or CD45.1$^+$CD4$^+$CD25$^+$ T cells from chimeric NOD mice ($0.5 \times 10^6$ per mouse) were simultaneously injected into NRG mice. Non-fasting blood glucose levels of the recipients were then monitored weekly for diabetes induction and mice were considered diabetic when two consecutive blood glucose readings were over 300 mg/dL. (**b**) Kaplan-Meier curve showing diabetes free survival. (**c**) Hematoxylin and eosin staining of pancreas section showing reduced T cell infiltration in T cell receiving NRG mice which also received host-derived CD4$^+$CD25$^+$ T cells. Magnification: 40×, Scale bar: 100 $\mu$m. NOD: non-obese diabetic, WT: wildtype.

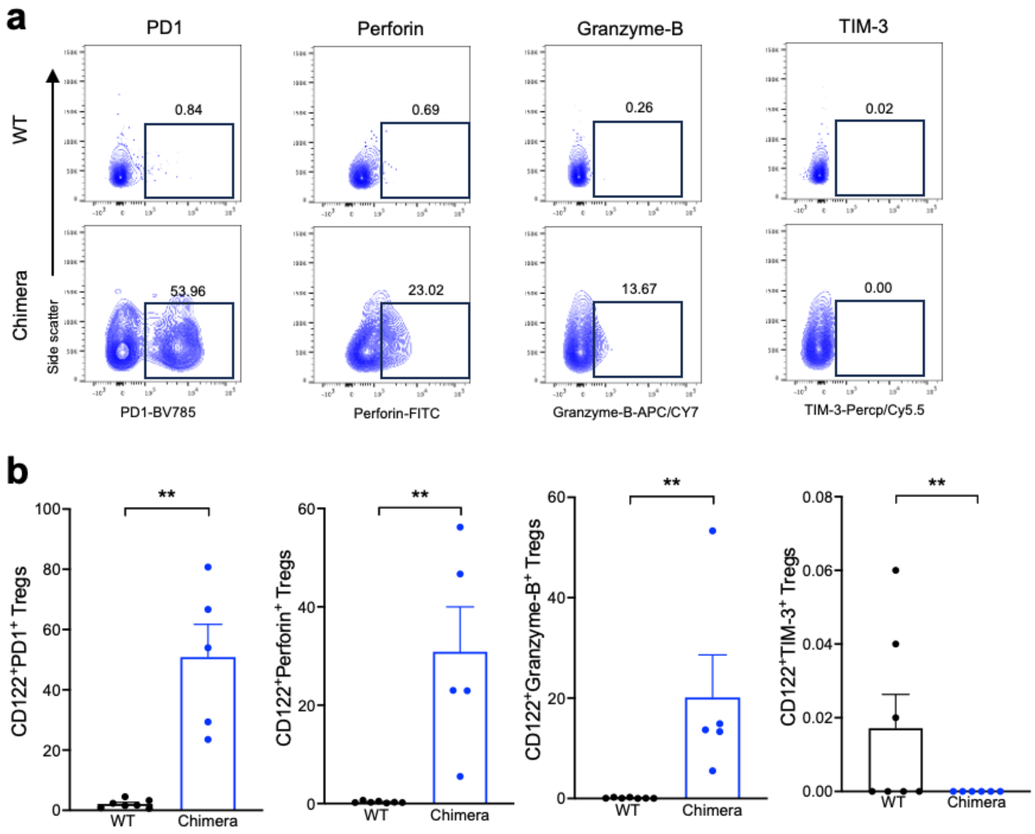


**Fig S7.** Phenotype of bone marrow-derived d-CD8$^+$CD122$^+$ Tregs. NOD chimeras were euthanized at day 60 post-HSCT and bone marrow cells were collected by flushing the femur with PBS. Age-matched WT C57BL/6 mice were used as controls. Dead cells were excluded by staining with Zombie Aqua™ Fixable Viability Kit. Cells were stained using antibodies against CD45.1, CD45.2, CD3, CD4, CD8, CD44, CD122, PD1, Perforin, Granzyme-B, and TIM-3. (**a**) Representative flow cytometry plots showing expression of PD1, Perforin, Granzyme-B, and TIM-3 in bone marrow-derived d-CD8$^+$CD122$^+$ Tregs from NOD chimeras and WT C57BL/6 mice. (**b**) Quantification of the percentage of PD1, Perforin, and Granzyme-B, and TIM-3 positive cells among CD8$^+$CD122$^+$ Tregs. **$p$<0.01. CD122$^+$ Tregs: CD8$^+$CD122$^+$ Tregs, PD1: programmed cell-death protein 1, TIM-3: T-cell immunoglobulin and mucin-domain containing-3, WT: wildtype.

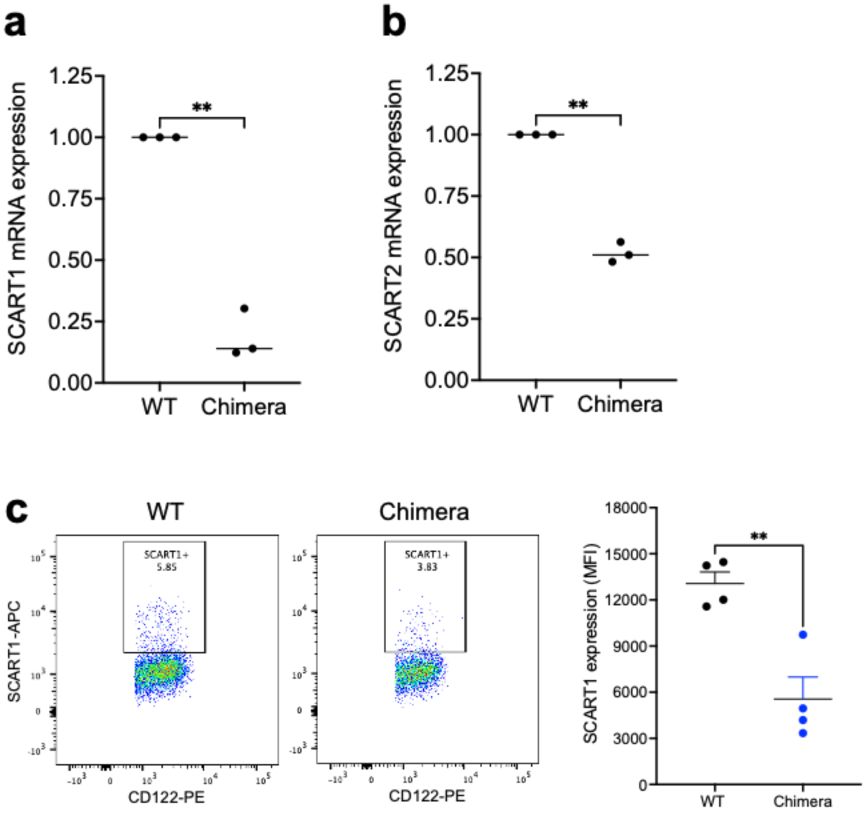


**Fig S8.** Donor-derived CD8$^+$CD122$^+$ Tregs have reduced expression Scart1 and Scart2. (**a-b**) qRT-PCR analysis was performed to verify the expression of the mRNA levels of Scart1 and Scart2. (**a**) mRNA expression of Scart1 in CD8$^+$CD122$^+$ Tregs. (**b**) mRNA expression of Scart2 in CD8$^+$CD122$^+$ Tregs. (**c-d**) Flow cytometric validation of Scart1 protein expression in CD8$^+$CD122$^+$ Tregs. Primary antibody against Scart1/CD163L1 (Novus Biologicals) was conjugated with FITC using a FlexAble FITC Plus Antibody Labeling Kit (Proteintech), following manufacturer's instructions. Lymphocytes from WT C57BL/6 and chimeric mice were then stained with antibodies against CD45.1, CD45.2, CD3, CD4, CD8, CD44, CD122, and CD163L1-FITC. (**c**) Representative flow cytometry plots showing expression of Scart1 protein in CD8$^+$CD122$^+$ Tregs. (**d**) Quantification of mean fluorescence intensity of Scart1 in CD8$^+$CD122$^+$ Tregs. ***p*<0.01. SCART: scavenger receptor family member expressed on T cells, WT: wildtype.

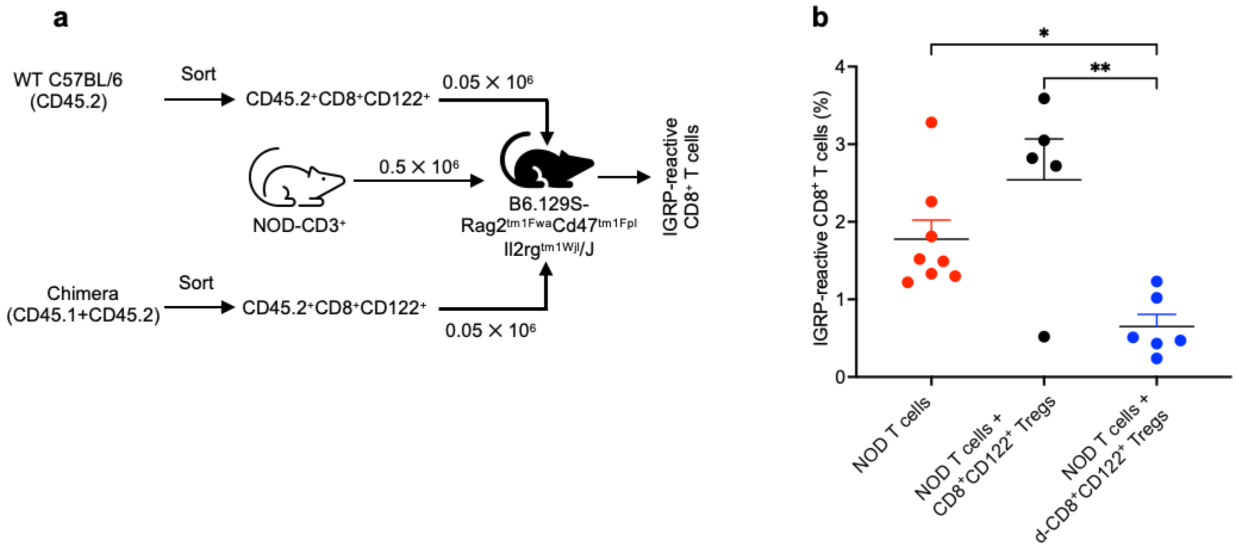


**Fig S9.** (**a**) Design of *in vivo* adoptive T cell transfer experiment. NOD chimeras were sacrificed at day 60 post-HSCT and d-$CD8^+CD122^+$ Tregs were sorted. A total of $0.5 \times 10^6$ $CD3^+$ T cells from WT NOD and $0.05 \times 10^6$ d-$CD8^+CD122^+$ Tregs were simultaneously transferred into Triple KO C57BL/6 mice. Group receiving $CD8^+CD122^+$ Tregs from WT C57BL/6 mice was used as control. (**b**) Frequency of IGRP-reactive $CD8^+$ T cells in Triple KO C57BL/6 mice at week 3 post T cell transfer. *$p$<0.05, **$p$<0.01. IGRP: islet-specific glucose-6-phosphatase subunit-related protein, WT: wildtype.

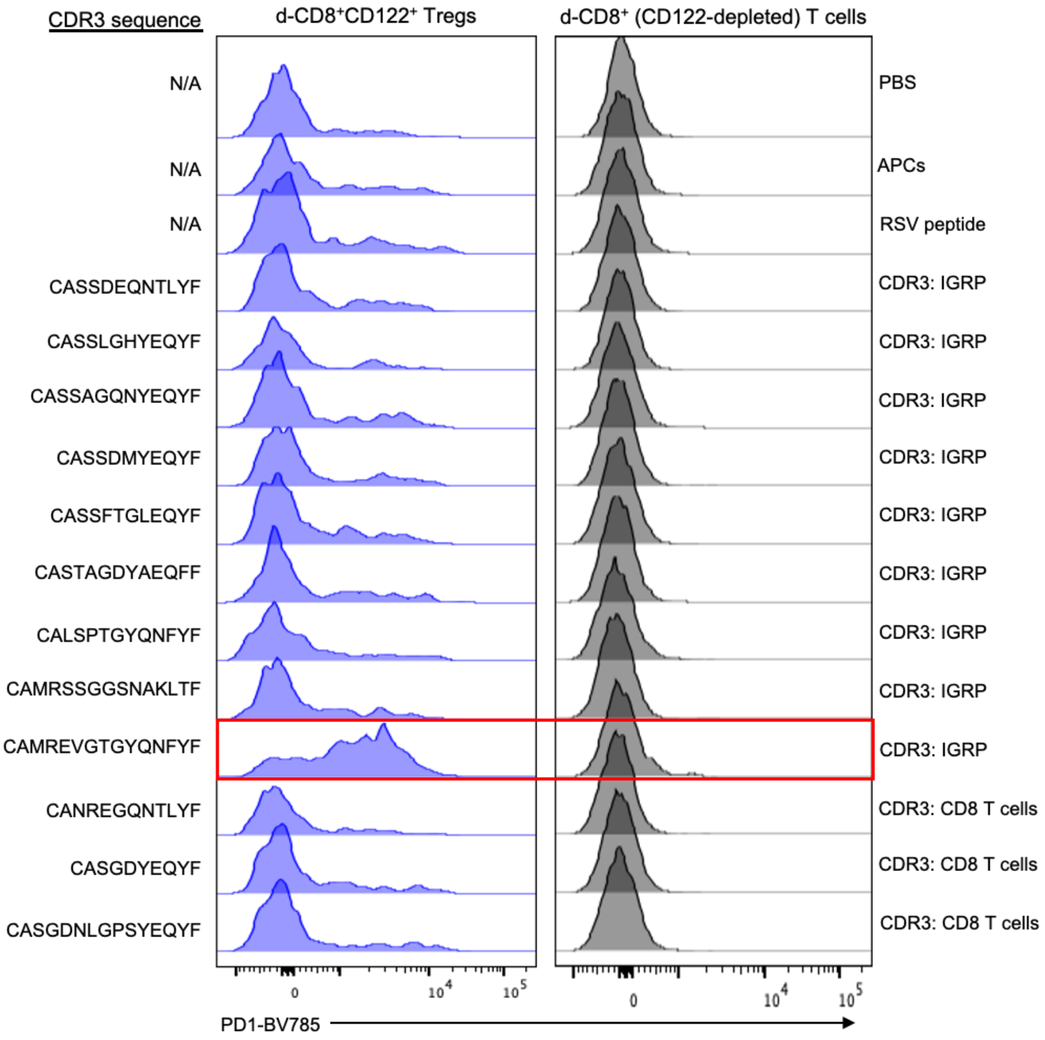


**Fig S10.** Screening of CDR3 peptides using activation assay. Nine CDR3 peptide sequences derived from IGRP-reactive $CD8^+$ T cells and three CDR3 peptide sequences derived from NOD $CD8^+$ bulk T cells were added separately in the co-culture of d-$CD8^+CD122^+$ Tregs or CD122-depleted d-$CD8^+$ T cells, and $CD11b^+$ monocytes as APCs for 48 h. RSV peptide (sequence: SYIGSINNI) was used as an irrelevant control. All peptides were used at a concentration of 20 μM. Representative histograms showing expression of PD1 in d-$CD8^+CD122^+$ Tregs (left) and CD122-depleted d-$CD8^+$ T (right) are shown. CDR3: complementarity-determining region-3, IGRP: islet-specific glucose-6-phosphatase subunit-related protein, N/A: not applicable, WT: wildtype.

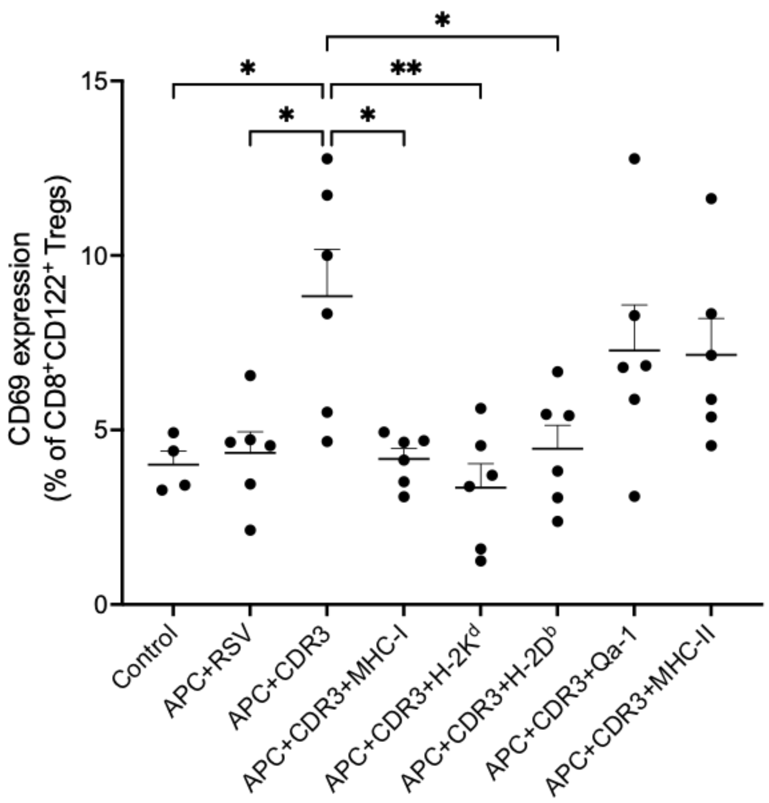


**Fig S11.** Blockade of MHC-I inhibits CDR3-driven activation of $CD8^{+}CD122^{+}$ Tregs. Pan MHC-I (BioXcell; BE0077), H-$2K^{d}$ (BioXcell; BE0104), H-$2D^{b}$ (BioXcell; BE0451), Qa-1 (BioXcell; BE0165), or MHC-II (BioXcell; BE0108) blocking antibodies at a concentration of 10 µg/mL were added into the coculture of d-$CD8^{+}CD122^{+}$ Tregs and $CD11b^{+}$ monocytes and 20 µM of CDR3 peptides. Percentage of $CD69^{+}$ d-$CD38^{+}CD122^{+}$ Tregs was quantified. Data were pooled from two independent experiments. *$p$<0.05, **$p$<0.01. APC; antigen presenting cell, CDR3; complementarity-determining region 3, MHC; major histocompatibility complex, RSV; respiratory syncytial virus.

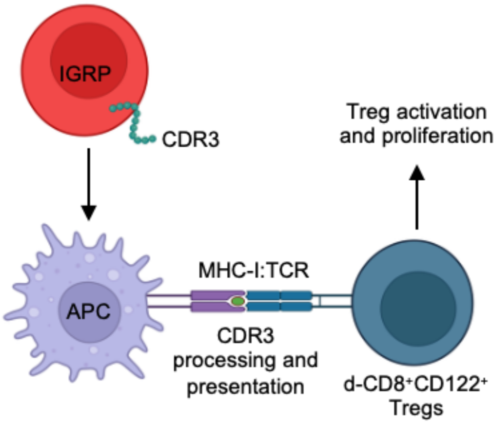


**Fig S12.** Proposed mechanism of IGRP-reactive CD8$^+$ T cell killing by d-CD8$^+$CD122$^+$ Tregs *in vitro*. The IGRP-reactive CD8$^+$ T cell-derived CDR3 peptides are processed by APCs and presented to d-CD8$^+$CD122$^+$ Tregs in the context of MHC-I. This leads to activation and proliferation of the d-CD8$^+$CD122$^+$ Tregs *in vitro*. APC: antigen presenting cell, CDR3: complementarity-determining region-3, IGRP: islet-specific glucose-6-phosphatase subunit-related protein, MHC-I: major histocompatibility complex I, TCR: T cell receptor.

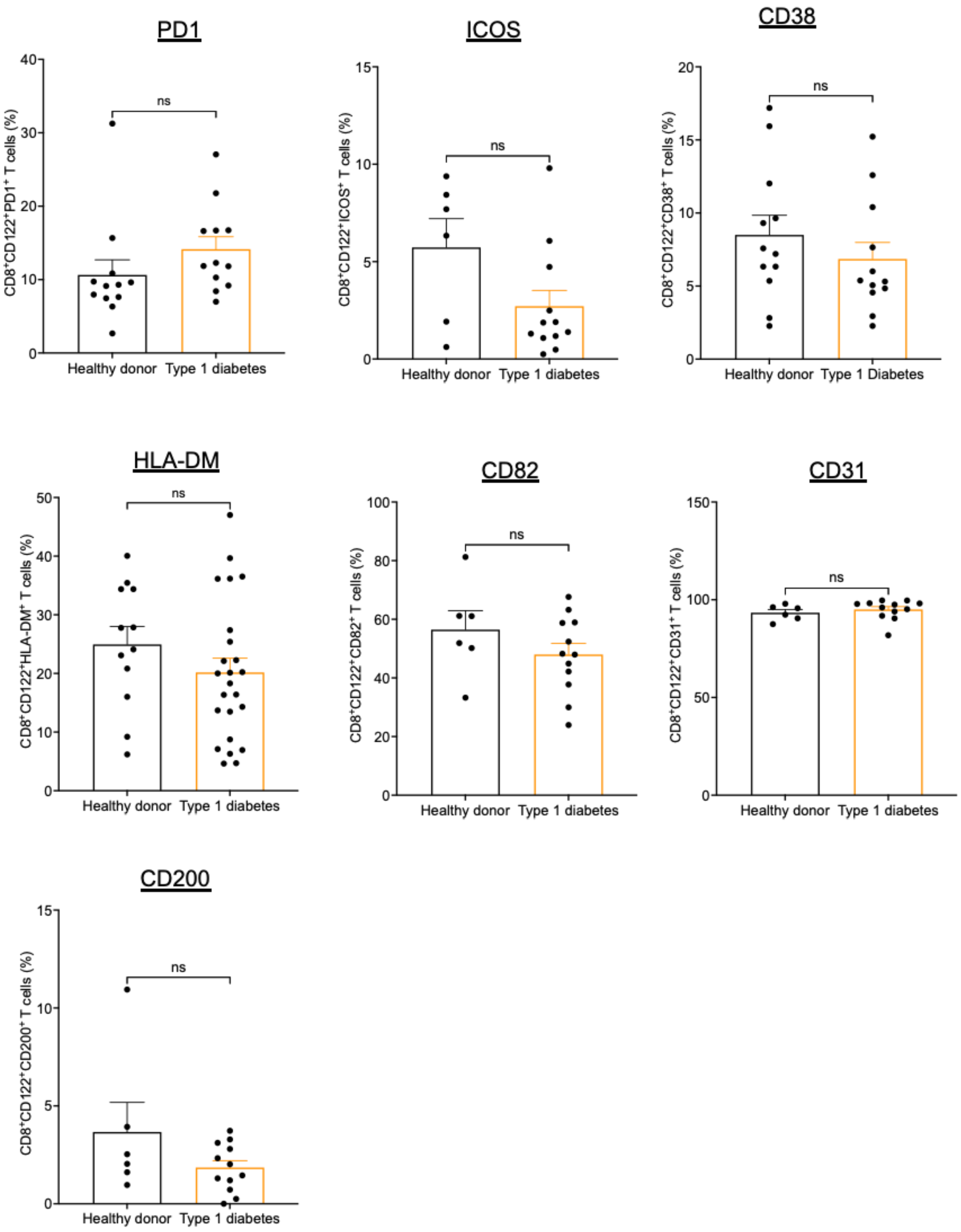


**Fig S13.** Phenotype of $CD8^+CD122^+$ Tregs in individuals with T1D. PBMC samples from T1D and HD controls were stained with antibodies against PD1, ICOS, CD38, HLA-DM, CD82, CD31, and CD200. Data represent mean ± SEM of two independent experiments. No statistically significant difference was observed in the expression of the surface markers among the $CD8^+CD122^+$ Tregs in T1D and HD controls. ns: not significant. ICOS: inducible T cell stimulator, HLA-DM: human leukocyte antigen-DM, PD1: programmed cell-death protein 1.

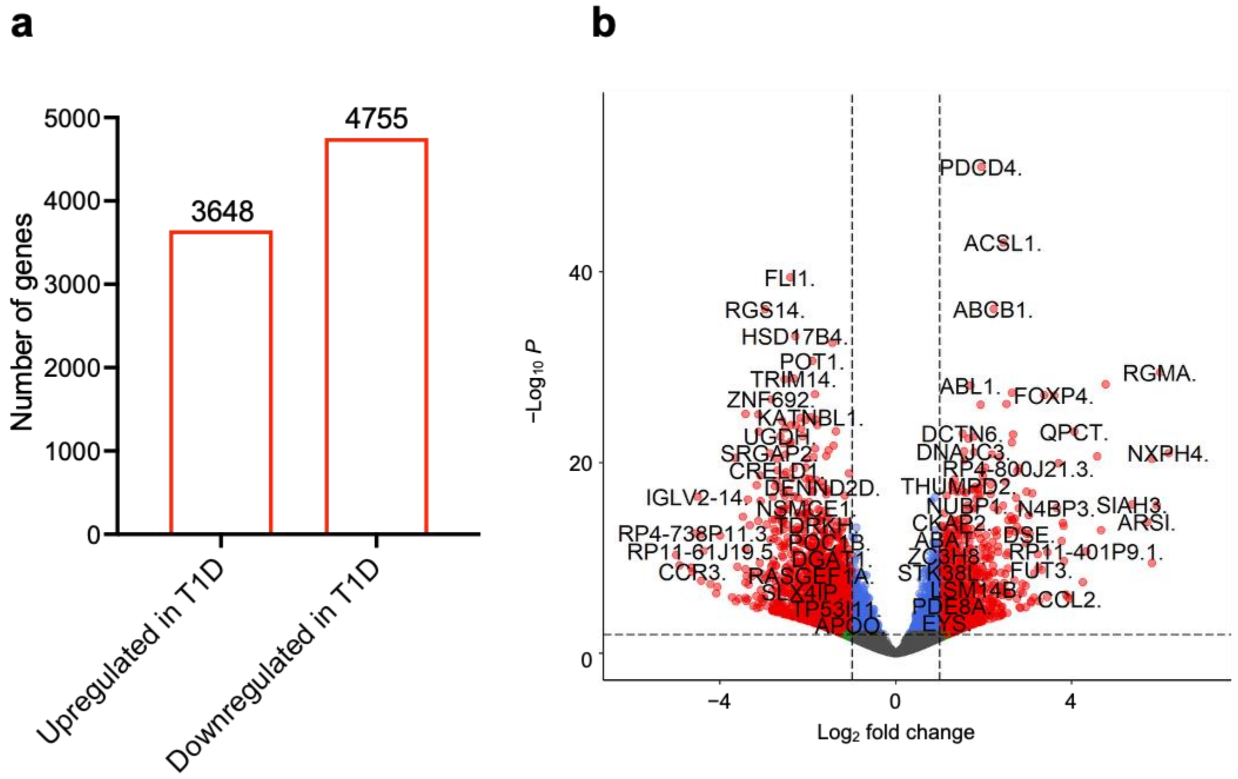


**Fig S14**. Bulk RNA sequencing of CD8$^+$CD122$^+$ Tregs (**a**) Total number of upregulated and downregulated genes in CD8$^+$CD122$^+$ Tregs from T1D *versus* HD with $p_{adj}$ <0.1. (**b**) Volcano plot showing gene expression $\log_2$ fold change and $-\log_{10}(P_{adj})$ in CD8$^+$CD122$^+$ Tregs from T1D *versus* HD. Significant differentially expressed genes are marked in red and are defined by a $\log_2$ fold change > |1| and a $P_{adj}$ < 0.001.

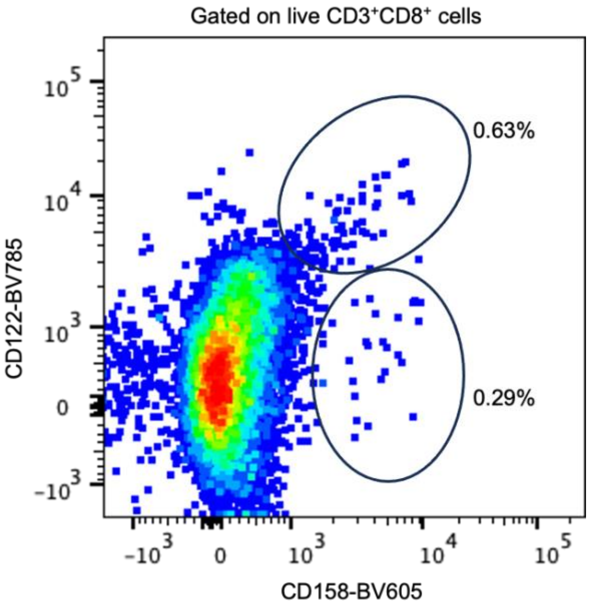


**Fig S15.** Representative flow cytometry plot showing overlapping phenotypic characteristics between $CD8^+CD122^+$ Tregs and $CD8^+CD158^+$ Tregs in human sample.

**Table S1.** Demographics of human samples used for Bulk RNA sequencing

| SN | Group | Sex | Age (years) | T1D onset duration |
|---|---|---|---|---|
| 1 | Healthy | unknown | unknown | N/A |
| 2 | Healthy | unknown | unknown | N/A |
| 3 | Healthy | unknown | unknown | N/A |
| 4 | Healthy | unknown | unknown | N/A |
| 5 | Healthy | unknown | unknown | N/A |
| 6 | Healthy | unknown | unknown | N/A |
| 7 | T1D | M | 17 | >1 year |
| 8 | T1D | F | 78 | >17 years |
| 9 | T1D | M | 43 | >26 years |
| 10 | T1D | M | 43 | >1 year |
| 11 | T1D | F | 73 | >30 years |
